# Real age prediction from the transcriptome with RAPToR

**DOI:** 10.1101/2021.09.07.459270

**Authors:** Romain Bulteau, Mirko Francesconi

**Affiliations:** Laboratoire de Biologie et Modélisation de la Cellule, Université de Lyon, ENS, UCBL, CNRS, INSERM, UMR5239, U 1210, F-69364 Lyon, France

## Abstract

Transcriptomic data is often affected by uncontrolled variation among samples that can obscure and confound the effects of interest. This is frequently due to unintended differences in developmental stages between samples. The transcriptome itself can be used to estimate developmental progression, but existing methods require many samples and do not estimate a real developmental time.

Here we present RAPToR, a simple and precise computational method that estimates the real age of a sample from its transcriptome, exploiting existing time-series data as reference. RAPToR works with whole animal, dissected tissue and single-cell data, for the most common animal models, humans and even for nonmodel organisms lacking reference data. We show RAPToR estimated age improves differential expression analysis by recovering the signal of interest when confounded with age. RAPToR will be especially useful in large scale single organism profiling because it eliminates the need for accurate staging or synchronization before profiling.

## Introduction

Genome-wide gene expression profiling is a powerful technique that provides a global and unbiased view of the transcriptional state of a biological system. However, the analysis of gene expression data can be complicated by uncontrolled and unknown sources of variance – which may be technical but also biological in nature^1^ – that can mask or confound the effects of variables of interest.

To tackle this problem, several methods have been developed to learn and remove hidden covariates (or surrogate variables) from the data, such as *Remove Unwanted Variance* (RUV)^2^, *Surrogate Variable Analysis* (SVA)^3^, or *Probabilistic Estimation of Expression Residuals* (PEER)4. However, a drawback of these methods is that the sources of variance usually remain obscure, therefore potentially interesting biological variance might also be removed.

A major source of variance in gene expression data of developing systems is often due to unintended differences in developmental progression across biological replicates or experimental conditions (Fig. 1a) that can confound (Fig. 1b) or mask (Fig. 1c) the effect of the variable of interest. This is especially true in organisms with rapid life cycles and highly variable growth speed such as worms, fruit fly or zebrafish, where numerous factors like genetic background, temperature, diet, crowding^5–9^, or even the physiological state of the previous generation^9^ substantially impact developmental speed. Carefully controlling for all conditions influencing development is therefore particularly challenging, but failing to do so can strongly impact gene expression. For example, in *C. elegans* even a few hours of development may result in 10,000 differentially expressed genes^10^. Hence, it is not surprising that around 50% of gene expression variance in the profiling of a large panel of *C. elegans* recombinant inbred lines^11^ is due to unintended developmental variation^12^ and that almost 38% of the datasets that did not intend to include development in a *C. elegans* gene expression database^13^ show substantial developmental variation in gene expression^10^.

**Figure 1.**
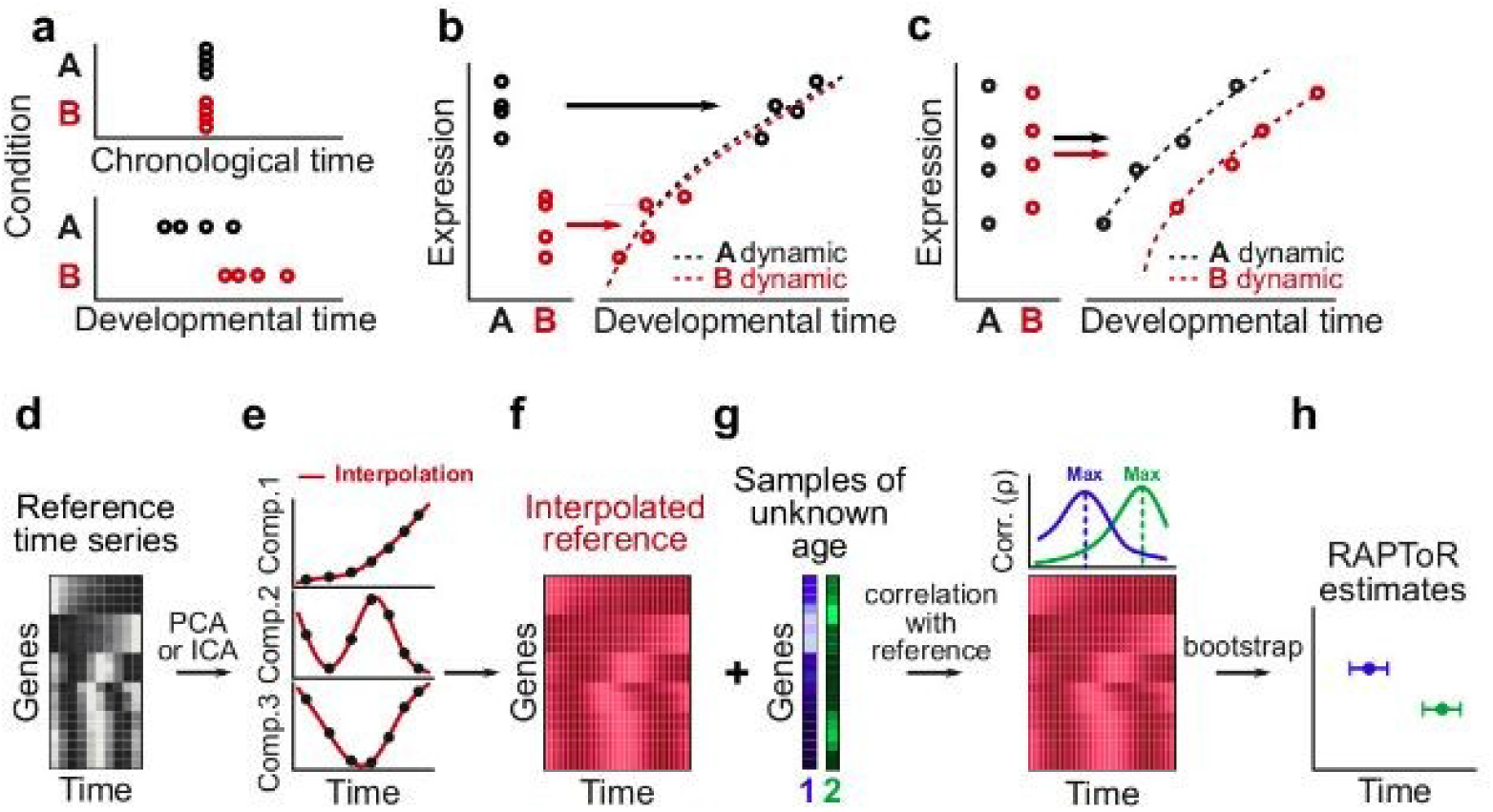
Estimate age from transcriptome using RAPToR. **a**, Cartoon of individuals sampled at identical chronological times in two conditions, resulting in different developmental age between groups due to the condition impacting developmental speed. **b-c**, Cartoons of differential expression analysis situations where hidden developmental variation is either (**b**) misinterpreted as an effect of the condition due development and condition being confounded, or (**c**) masking an effect of the condition due to developmental spread. **d-f**, RAPToR staging exploits existing reference time-series expression data (**d**). This data is first decomposed into principal / independent components which are interpolated with respect to time (**e**). Interpolated reference is then reconstructed at gene level with interpolated components and gene loadings (**f**). **g-h**, For each sample, a correlation profile is built by computing genome-wide Spearman correlation with every time point of the reference (**g**). The reference time with maximal correlation becomes the estimate, and bootstrapping on random gene subsets defines a confidence interval (see methods) (**h**).

Estimating the real physiological age of the samples and identifying hidden developmental variation between them is important first to quantify the impact of the perturbation of interest on developmental speed; second, to distinguish perturbation-specific from unspecific gene expression changes caused by development; third, to uncover time-specific effects of the perturbations under study^12^ by including inferred age as a covariate in expression data analyses (such as differential expression analysis). In yeast, analogous ideas successfully identified genetic and environmental perturbations impacting specific phases of the cell cycle^14^ and direct and specific effects of 700 gene deletions on gene expression after removing the main source of variance (25%): a shared expression signature of cell cycle and growth rate^15^.

Extracting developmental progression from transcriptomes has recently become a topic of intense research, especially after the advent of single-cell RNASeq. Many algorithms have been developed that learn developmental progression from large scale bulk, single cell, or whole-organism transcriptomic data and sort samples along those trajectories (e.g. Slingshot^16^, DPT^17^, Monocle^18^, BLIND^19^). However, a major drawback of these trajectory-learning algorithms is that they require large amounts of samples to learn the trajectory of developmental expression changes directly from the data. Moreover, they only output dataset-specific ranks or arbitrary values usually referred to as “pseudo-times”, making it difficult to compare results across datasets or conditions.

To overcome these limitations, we developed RAPToR (**R**eal **A**ge **P**rediction from **T**ranscriptome staging **o**n **R**eference), a computational framework that exploits available time-series gene expression data as reference to determine the absolute developmental age of even a single sample from its transcriptome with high precision. We implemented RAPToR in R (available at https://github.com/LBMC/RAPToR), providing references to stage *C. elegans, D. melanogaster, D. rerio*, and *M. musculus* development from gene expression.

We show that RAPToR successfully estimates age during development and aging, estimates tissue-specific developmental age from whole-organism data, and can also estimate age of one species using another species as reference. Finally, we show how to use estimated ages to quantify a perturbation effect on developmental speed, and recover perturbation-specific effects on gene expression even when the perturbation is completely confounded by development.

## Results

### RAPToR Design

We set out to develop a strategy to continuously stage development from gene expression that would be effective even for experiments with a limited number of samples, where trajectory learning methods are not applicable. We reasoned that we could exploit existing developmental time-series data as reference to stage samples individually by taking the time point of the reference that has maximum correlation with a sample transcriptome as the age estimate. In this way, the age of each sample is inferred independently from others and outliers only influence their own staging (as opposed to trajectory-based approaches). Furthermore, age estimates acquired on the same reference should be comparable even across different experiments, conditions and genetic backgrounds.

However, using time of maximum reference correlation limits the precision of age estimates to the temporal resolution of the reference. To overcome this limitation, we interpolate reference gene expression (Fig. 1d) with respect to time in a dimensionally reduced space (Fig. 1e, Sup. Note 1), generating interpolated expression profiles potentially at any time between the original reference time points (Fig. 1f).

The sample age estimate is simply the time point of maximum Spearman correlation between the interpolated reference and the sample gene expression (Fig. 1g). We then compute a confidence interval of the estimate by bootstrapping on genes (Fig. 1h, methods).

We implemented this strategy in RAPToR, an R package where we provide functions to interpolate references and stage samples.

### RAPToR accurately estimates the developmental age of common model organisms

To test RAPToR in the most commonly used animal model organisms, we built interpolated references exploiting existing time-series data on *C. elegans* roundworm embryonic and larval development^20–22^, zebrafi*s*h embryonic and larval development^23^, mouse^24^, and fruit fly^25^ embryonic development (Sup. Table 1) and then staged independent time-series experiments of *C. elegans* late-larval development^26^ and zebrafish^27,28^, mouse^29^, and fruit fly^27^ embryonic development.

We found RAPToR age estimates accurately match chronological age for both *C. elegans* and zebrafish (R^2^>0.99, Fig. 2a, 2b), and morphological staging (somite number) for mice (R^2^=0.95, Fig. 2c). Age estimates of fly single-embryos27 less accurately match chronological age especially at later stages (R^2^=0.74, Fig. 2d). However, this is likely due to the single individual nature of data as any inter-individual variation in developmental speed would not be averaged out as in bulk data. Indeed, the authors used BLIND^19^ – a trajectory-learning method to re-rank their samples^27^ similarly to RAPToR (ρ>0.99, Sup. Fig. 1). RAPToR estimates do in fact enhance expression dynamics captured by principal components (Fig. 2e, 2f, Sup. Fig. 1) and for the majority of genes (Sup. Fig. 1) compared to chronological age (see methods). Thus, RAPToR estimates the true physiological age of individuals and reveals the heterogeneity of their developmental speeds.

**Figure 2.**
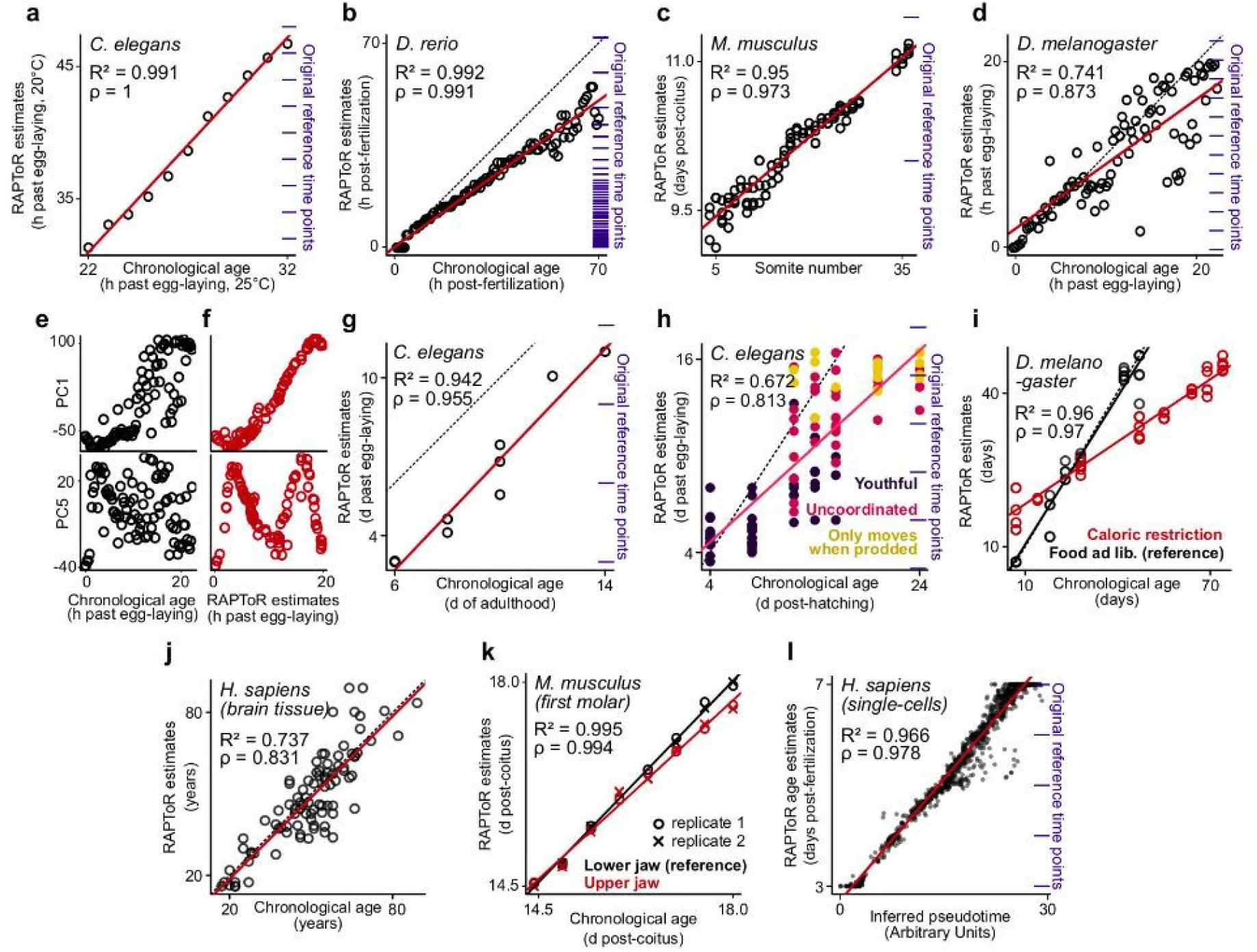
Evaluating RAPToR’s performance by staging time-series of model organisms. **a,b**, Chronological age vs. RAPToR estimates of *C. elegans* late-larval samples^26^ (linear model is y = −4.7 + 1.6x) (**a**), and *D. rerio* embryo samples^27^ (linear model is y = 0.7x) (**b**) **c**, Somite-number vs. RAPToR estimates of *M. musculus* embryo samples^29^ (linear model is y = 9.2 + 0.05x) **d**, Chronological age vs. RAPToR estimates of *D. melanogaster* embryo samples^27^ (linear model is y = 1.3 + 0.77x) **e, f**, Selected Principal Components of the data staged in (**d**), plotted in black along chronological age (**e**) and in red along RAPToR estimates (**f**). **g-j**, Chronological age vs. RAPToR estimates of adult *C. elegans* bulk samples^33^ (**g**) and single-worms^34^ (**h**), of adult *D. melanogaster*^35^ (**i**), and of human brain tissue^36^ (**j**). **k**, Chronological age vs. RAPToR estimates of dissected samples of upper jaw first molars from *M. musculus* embryos staged using the lower jaw samples as reference^37,38^. **l**, Inferred pseudotime vs. RAPToR estimates of *H. sapiens* embryo single cells^39^. **a-d,g,h**, Staged samples^26,27,29,27,33,34^ and references^20,23–25^ are from independent time-series experiments. Original time points of the references within the plot area are shown to the right (blue), but the references can span much longer coverage.

### Reference interpolation dramatically increases temporal resolution and accuracy of age estimates

Crucially, reference interpolation allows staging with an accuracy far beyond the original sampling resolution of reference time-series. Indeed, RAPToR accurately stages a dense zebrafish time course^28^ with over 40 times the temporal resolution of the reference before interpolation (Sup. Note 1, Sup. Fig. 2, methods). RAPToR estimates also stay remarkably accurate and precise even when staging samples with a few hundred genes, or with noisy data (Sup. Note 1, Sup. Fig. 3, 4, 5, 6) and are robust to reference interpolation parameters (Sup. Note 1, Sup. Fig. 7, 8, Sup. Table 2).

### RAPToR correctly infers developmental speed scaling factors

RAPToR estimates are relative to the reference chronological age. Thus, one can use RAPToR to stage samples with known chronological age to estimate developmental speed differences or scaling factors with a reference. For example, staging a *C. elegans* developmental time-series grown at 25°C^26^ on the reference grown at 20°C^20^ recapitulates the expected 1.5 fold increase in developmental speed due to temperature increase^20^ (Fig. 2a, Sup. Note 1).

### RAPToR performs well on aging

While RAPToR works very well with robust expression changes during development, aging and aging-related gene expression changes are widely known to be heterogeneous, stochastic, and strongly influenced by environmental factors^30–32^. This can potentially limit the applicability of RAPToR to aging. In fact, RAPToR performs poorly between independent aging time-series (but works within experiments, Sup. Note 1, Sup. Fig. 9) with references built using the whole transcriptome. We reasoned RAPToR performance could increase by strengthening the aging signal in the reference. Indeed, by building RAPToR references restricting to genes with robust monotonous trends along aging (see methods) we could successfully estimate aging in *C. elegans* bulk^33^ (R^2^=0.94, Fig. 2g) and single-worm^34^ samples (R^2^=0.67, Fig. 2h), *Drosophila*^35^ (R^2^=0.96, Fig 2i), and humans^36^ (R^2^=0.74, Fig.2j, Sup. Note 1, Sup. Fig. 10). Importantly, single-worms staged older than their chronological age behaved like older individuals^34^ and vice versa (Fig. 2h), moreover age estimates of flies under caloric restriction are consistent with the expected lifespan extension^35^ (Fig. 2j). This shows that RAPToR age estimates recapitulate true differences in biological age across individuals or environmental conditions. We conclude that RAPToR reliably infers aging from transcriptomic data.

### RAPToR stages dissected tissue samples well

We tested RAPToR on expression profiling from dissected tissues – where variation in cell type composition and relative amount might potentially confound staging – using time-series of *M. musculus* upper and lower-jaw first molar development^37,38^. Since these two organs have very similar development^37^, we built a lower-jaw reference to stage the upper-jaw samples (see methods). RAPToR not only accurately estimates age (R^2^>0.99, Fig. 2k), but also correctly infers the known developmental delay of upper molars compared to lower molars^37,38^. Thus, despite potential confounders, RAPToR is effective and precise on dissected tissue samples.

### RAPToR accurately stages single-cell data

Single-cell expression data is usually much sparser and noisier than bulk data, which might potentially limit RAPToR performance. We therefore tested whether RAPToR could stage human early-embryo development single cells^39^ by building a reference with a random subset of the data and staging the remaining cells (see methods). Age estimates not only match the chronological time (R^2^=0.87, Sup. Fig. 11), but also strongly correlate with the pseudo-times computed by the authors^39^ (R^2^=0.97, Fig 2l). Therefore, despite the sparsity of the data, RAPToR ranks cells as well as pseudo-time methods specifically designed for single-cell data but at the same time provides a real biological time for each cell.

### RAPToR age estimates are robust to genetic variation in gene expression

Variable genetic background is another potential confounder, so we tested RAPToR performance on expression data for over 200 *C. elegans* recombinant inbred lines (RILs) showing extensive genetic variation in gene expression^11^. RAPToR closely matches previous estimates from a trajectory-learning approach^13^ (R^2^=0.94, Sup. Fig. 12), thus confirming the RILs span mid-larval to young adult stage, a period with vast expression changes both in the soma (molting) and the germline (spermatogenesis, oogenesis) of worms.

We noticed that some gene expression dynamics in the RILs are advanced and others delayed compared to the reference (Fig. 3a, 3b, Sup. Fig. 13, Sup. Note 1). Shifts between soma and germline development (soma-germline heterochrony) are easily induced by environmental and physiological changes in *C. elegans*^9,40^. Indeed, a consistent enrichment of soma and germline genes in advanced and delayed dynamics respectively suggests soma-germline heterochrony between the reference and the RILs (Sup. Fig. 13).

**Figure 3.**
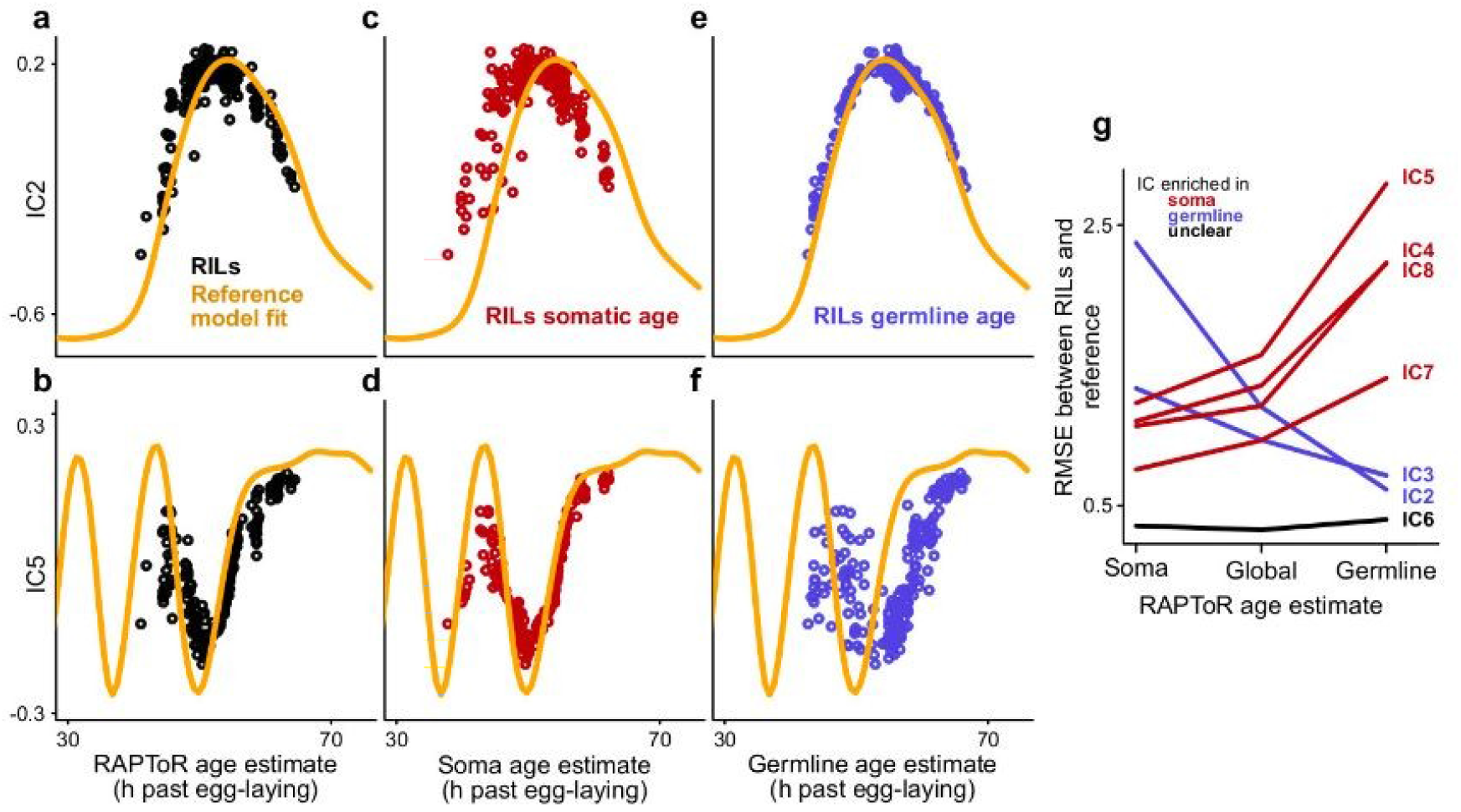
Tissue-specific staging. **a-b**, Selected Independent components from ICA on joint *C. elegans* Recombinant Inbred Lines^11^ (RILs, dots) and reference data^21^ they were staged on (orange line). **c-f**, As in (**a-b**), with RILs plotted in red along soma age (**c-d**), and in blue along germline age (**e-f**). **g**, RMSE between RILs and reference for independent components 2-8 when using soma, global or germline age estimates.

### Tissue specific staging enables quantification of heterochrony

To confirm this we used germline- and soma-specific gene sets^22,26^ to separately stage the germline and soma of the RILs (see methods, Sup. Fig. 12). We find germline- and soma-specific dynamics align better on the reference when staged with the corresponding gene set (Fig. 3d, 3e, 3g) while they are otherwise shifted (Fig. 3c, 3f), confirming heterochrony between reference and RILs. Thus tissue-specific staging outperforms global staging in case of heterochrony between the reference and the samples to stage.

Beyond differences between the reference and RILs, we noticed that tissue-specific staging also decreases variance among the RILs. Indeed, germline genes are better fit by germline than soma age and vice versa, suggesting soma-germline heterochrony among the RILs (Sup. Fig. 14). However, when searching for the genetic basis of this heterochrony with a multivariate QTL analysis, we found no significant genetic locus (even at an FDR of 0.5) and overall no significant amount of genetic variance in heterochrony (Sup. Note 1), which is therefore likely due to unknown and uncontrolled environmental differences or to a very complex genetic architecture not captured by the model.

In summary, by using tissue-specific gene sets RAPToR provides accurate tissue-specific age estimates from whole-organism expression despite varying genetic background.

### Staging on references of a different species

Developmental time-series data are often unavailable for non-model organisms. However, gene expression dynamics during development are often well-conserved across related species, especially during the phylotypic stage^41^. Seeing the robustness of RAPToR to genetic variation within species, we decided to test how well RAPToR can stage one species on a related species.

Staging time-series of embryo development across 6 *Drosophila* species^41^ on a *D. melanogaster* reference using orthologs indeed results in accurate age estimates (R^2^>0.99, Fig. 4a) despite decreasing overall correlation with increasing phylogenetic distance (Fig. 4b). Moreover, we infer between-species growth speed factors matching those found by the authors (Sup. Table 3). Importantly, we also detect small age differences between replicates of each time point, which refine expression dynamics (Sup. Fig. 15), thus reducing noise in the data (Sup. Fig. 16).

**Figure 4.**
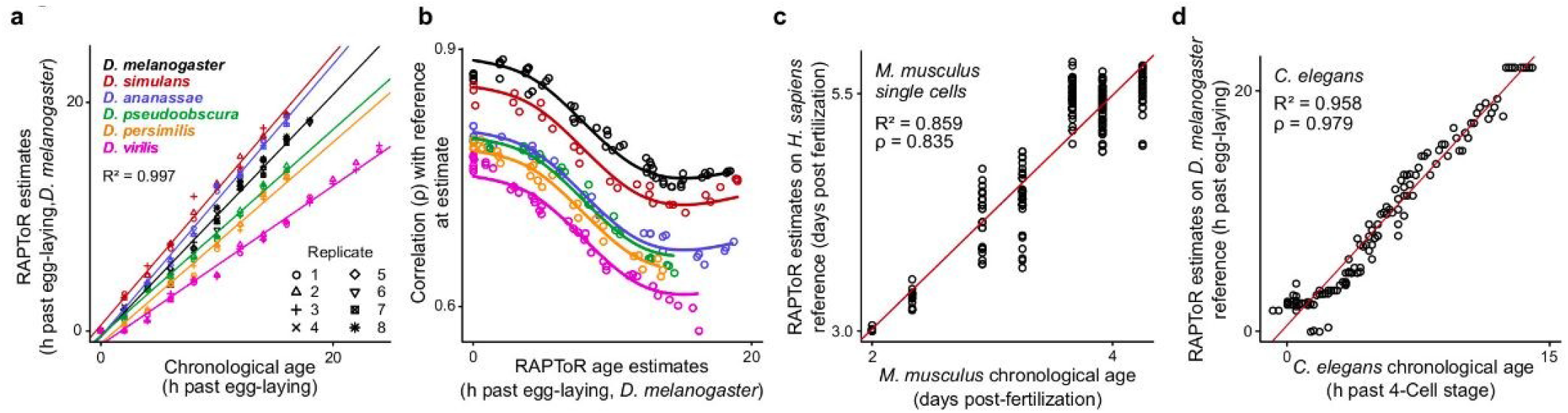
Staging samples cross-species. **a**, Chronological age vs. RAPToR estimates for time-series of embryo development of 6 *Drosophila* species^41^ staged on a *D. melanogaster* reference^25^ (see also Sup. Fig. 15). **b**, Spearman correlation between samples from (**a**) and the reference at age estimate, along RAPToR estimates. **c**, Chronological age vs. RAPToR estimates for single-cells of *M. musculus* embryos^44^, staged on a *H. sapiens* single-cell reference^39^ using orthologs. **d**, Chronological age vs. RAPToR estimates for *C. elegans* embryo samples^27^, staged on a *D. melanogaster* reference^25^ using orthologs.

Encouraged by this, we probed RAPToR limits by staging samples on more distant reference species. We were able to stage human^42^ on cow^43^ whole-embryos (R^2^ = 0.83, Sup. Fig. 17), as well as early-embryo mouse single-cells^44^ on a human reference^39^, matching both chronological age (R^2^ = 0.86, Fig. 4c) and pseudo-times (rho=0.95, Sup. Fig. 18). To our surprise, we could even successfully stage *C. elegans* embryogenesis^27^ on a *D. melanogaster* reference (R^2^ = 0.96, Fig. 4d, Sup. Note 1, Sup. Fig. 19), two species separated by 600 million years of evolution^45^.

Which biological processes with highly conserved dynamics during embryogenesis could account for this accurate staging? We found that gene expression signatures of decreasing cell proliferation shared across phyla^27^ and signatures of muscle development or cell differentiation are necessary and almost sufficient for accurate staging (Sup. Note 1, Sup. Fig. 19, 18, Sup. Tables 4-11, methods).

Thus RAPToR can stage non-model organisms using available close species data and perform well even in extremely distant species, when applied to developmental stages with highly conserved developmental dynamics.

To summarize, RAPToR performs well across the organisms, sample types, and diverging genetic backgrounds and species we tested, yielding estimates that are precise, accurate thanks to interpolation, and robust to gene set size changes.

### RAPToR finds hidden drug effects on germline development

RAPToR absolute age estimates are useful in many ways. First, rather than just getting a list of differentially expressed genes from profiling data, RAPToR precisely quantifies the effect of perturbations on developmental timing, including in a tissue-specific way. For example, tissue-specific staging of *C. elegans* exposed to three concentrations of mefloquine, dichlorvos, and fenamiphos^46^ found that all three drugs induce a similar germline-specific and dose-dependent developmental delay (Fig. 5a, Sup. Note 2, Sup. Fig. 20).

**Figure 5.**
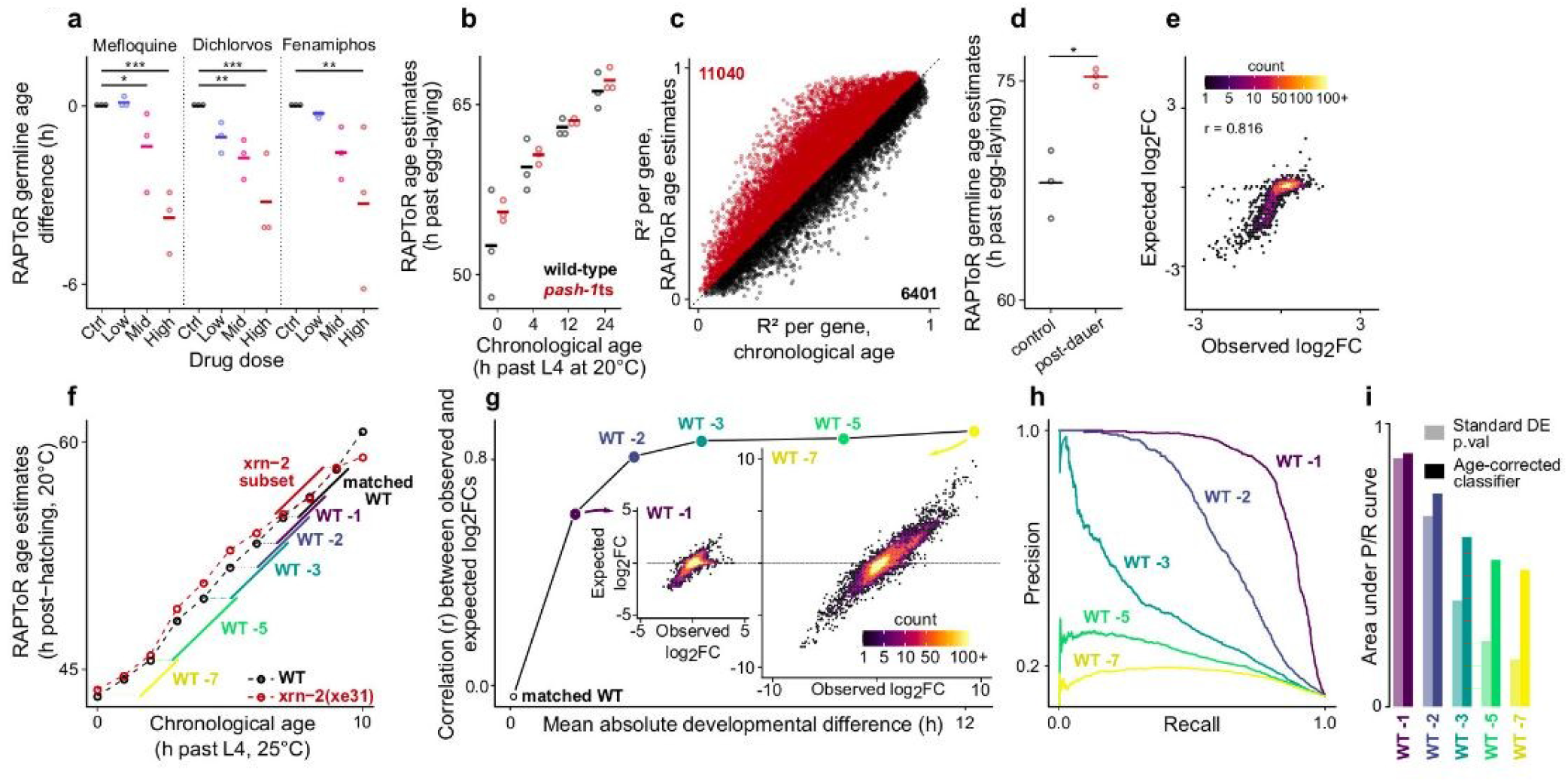
Quantifying and correcting for developmental effects using RAPToR age estimates. **a,** Effect of increasing drug dose exposure^46^ on RAPToR estimates of *C. elegans* germline age (see methods, Sup. Fig. 20). **b,** RAPToR estimates vs. reported chronological age highlight large developmental spread within time points of *C. elegans* WT and *pash-1ts* time-series^47^ (see Sup. Note 2, Sup. Fig. 21). **c**, R^2^ per gene of identical models with chronological age, or RAPToR estimates. Genes and gene counts above and below the dashed line (x=y) are indicated in red and black respectively. **d,** Germline age estimates of control and PD *C. elegans* adults^48^. **e**, Germline genes logFCs between control and PD from (**d**) compared to logFCs expected from developmental time difference only (see Sup. Note 2, Sup. Fig. 24). **f,** Chronological age vs. RAPToR estimates of a time-course of *C. elegans* WT and *xrn-2* late larval development^49^. Sample subsets defining a gold standard of truly DE genes and shifted WT sets used in subsequent panels are color-coded. **g**, Correlation of observed logFCs and expected developmental logFCs computed from the interpolated reference between the *xrn-2* subset and increasingly shifted WT sets from (**f**). (see Sup. Note 2). **h**, Precision-Recall curves showing the performance of a standard DE model p-value for each shifted WT subset in detecting gold-standard DE genes. **i**, Area under precision-recall curves (AUPRC) of standard DE model p-value (**h**) or of the age-corrected classifier for each shifted WT subset in detecting gold-standard DE genes (see Sup. Note 2). *: p < 0.05, **: p < 0.01, ***: p < 0.001. PD: post-dauer, WT: wild-type, logFC: log2 fold-change, DE: differentially expressed or differential expression, FDR: false discovery rate.

### RAPToR increases statistical power in differential expression analyses

Even when known chronological age is included as a model covariate in differential expression (DE) analyses, replacing it by RAPToR age estimates increases statistical power. For example, using RAPToR estimates instead of chronological age when analyzing expression changes in *C. elegans pash-1* vs. wildtype^47^ (Fig. 5b), detects up to 60% more DE genes in *pash-1* and 10% more DE genes across development thanks to overall better model fits (Fig. 5c, Sup. Fig. 21, Sup. Note 2).

### Quantifying differential expression due to differences in developmental progression

If an experimental condition strongly impacts developmental speed but perturbed and control samples are collected at the same chronological – and therefore different physiological – time (Fig. 1a), the variable of interest will be completely confounded with development. Thus, purely developmental expression changes are wrongly attributed to the perturbation of interest (Fig. 1b). As an example, we reanalyze a dataset comparing young adult *C. elegans* that developed through dauer state (post-dauer) to controls that did not^48^. The authors found a down-regulation of spermatogenesis-associated genes and an up-regulation of oogenesis-associated genes from which they concluded that post-dauer animals have reduced spermatogenesis and increased oogenesis. However, as *C. elegans* switch from sperm to egg production during development, this pattern could simply be explained by post-dauer samples being physiologically older than controls. This is indeed what RAPToR found (Fig. 5d, Sup. Fig. 22, Sup Note 2). Furthermore, strong correlation (r > 0.8) between the observed expression changes in germline genes and the expected developmental expression changes calculated from matching time points in the reference (Fig. 5e, Sup. Fig. 22, 23, Sup Note 2) suggests that, despite synchronization efforts, most of the initially observed DE is due to uncontrolled differences in developmental progression.

### Recovering specific effects even when the variable of interest is completely confounded by development

We reasoned that integrating RAPToR age estimates and developmental gene expression from the reference in the DE analysis should allow us to extract perturbation-specific expression changes even when the variable of interest is completely confounded with development (Sup. Note 2, Sup. Fig. 24). We tested this using a *C. elegans* larval development time-series of *xrn-2* mutant and relative wild-type (WT) control sampled every 1.5h^49^. We defined a gold standard of truly DE genes in the mutant, which allowed us to vary the age difference between mutant and WT and quantify: first the amount, intensity, and variance of expression changes due to development (Fig. 5f-g, Sup. Fig. 24, Sup. Note 2); second, the deleterious impact of these developmental expression changes on the performance of a standard analysis in detecting truly DE genes; third the improvement obtained by integrating RAPToR estimates and reference expression data in the model. As expected, with increasing age differences between mutant and WT, the performance of a standard DE test sharply decreases (Fig. 5h). However, performance is greatly recovered by integrating reference data in the model, especially for large age differences when the mutant effect would be fully confounded by development (Fig. 5i, Sup. Fig. 24, Sup. Note 2).

In summary, we showed that using RAPToR and reference data it is possible to measure the impact of development in gene expression analyses and recover the specific effect of a perturbation even when completely confounded with development.

## Discussion

We presented here RAPToR, a computational strategy to accurately estimate the age of samples from their gene expression profile. Unlike trajectory-based methods, RAPToR exploits existing reference time-series data to continuously stage each sample separately, providing several advantages: first, it eliminates the need for large datasets to infer developmental trajectories; second, it provides absolute developmental times that are comparable across data sets, conditions, genetic backgrounds, profiling technologies and other covariates; third, outliers have no impact on the staging of other samples as each sample is staged independently.

While RAPToR is limited by the existence of reference time-series data, interpolation allows precise staging well beyond the resolution of the original reference data, enabling the use of sparse time-series as references. RAPToR estimates age both during development or aging in most common animal models and humans, from bulk, single-individual, dissected tissue or single-cell expression profiles, and can also infer tissue-specific age from whole-organism profiles. Importantly, RAPToR can stage one species using a close species as reference, which dramatically expands the scope of RAPToR, including to non-model organisms. We showed how RAPToR absolute estimates can be exploited in many ways: to detect the effect of a perturbation on developmental speed; as model covariates to increase statistical power to detect differential expression; finally, we showed that even in the extreme scenario when the perturbation of interest is completely confounded with development, it is still possible to recover genuine perturbation-specific expression changes by integrating reference data in differential expression analysis.

We anticipate our strategy of staging post-profiling with RAPToR will be especially useful in large scale singleorganism profiling experiments since it eliminates the need for synchronization or for tedious and potentially difficult steps of accurate staging before profiling.

To conclude, we remark that our approach is not restricted to development but can in principle be applied to any process with robust underlying reference gene expression dynamics (e.g. cell differentiation, cell cycle, disease progression, drug response) and its scope will only expand with the increasing availability of timeseries profiling data.

## Methods

Analyses were all performed using the R statistical software (v3.6.3)

## Data accessibility

All the data used in this study were previously published and deposited in public databases or accessible by request to the authors. The data from Sémon et al. is, at the time of writing, awaiting publication^38^. The full list of datasets and accession numbers is given in Supplementary Table 12.

The code to download and (pre)process the data, perform the analyses and generate the figures of this paper can be found at https://gitbio.ens-lyon.fr/LBMC/qrg/raptor-analysis

## Data pre-processing

Probe or gene IDs of datasets were converted to standard IDs (WBGene IDs for *C. elegans*, FBgn IDs for *D. melanogaster*, Ensembl IDs for *D. rerio, M. musculus, H. sapiens* and *B. taurus*). When multiple probes or IDs matched a single standard ID, they were mean-aggregated for microarray, sum-aggregated for RNA-seq counts. IDs with no standard ID match were dropped.

For RNA-Seq datasets, gene-level TPM data was used when available, or computed from raw counts using transcript lengths from the Ensembl biomart (v99). No remapping of the transcriptomes was done, aside from the *M. musculus tooth* data (see below). No background correction was applied to microarray data.

Samples were considered of poor quality and discarded when the 99^th^ percentile of the distribution of their Spearman correlation coefficients with others samples fell below a threshold defined below for each dataset.

Expression values for all datasets were quantile-normalized using the *normalizeBetweenArrays* function from *limma*^50^ (v3.42.0) on ***log(X +1)*** transformed values unless otherwise specified.

## RAPToR implementation

Our method is implemented in an R package: RAPToR (v1.1.4), which can be downloaded and installed from the following url. https://github.com/LBMC/RAPToR

Functions for staging samples, plotting results, interpolation and building references are included in the package. Detailed vignettes on general usage, reference building and showcases are also provided with the package.

Auxiliary R data-packages include references for *C. elegans* (embryonic, larval and young adult to adult development, https://github.com/LBMC/wormRef), *D. melanogaster* (embryonic development, https://github.com/LBMC/drosoRef), *D. rerio,* (embryonic and larval development, https://github.com/LBMC/zebraRef) and *M. musculus* (embryonic development, https://github.com/LBMC/mouseRef).

### Reference interpolation

Let ***X*** (***m × n***) be the gene expression matrix of ***m*** genes by ***n*** samples. The matrix is first gene-centered such that ***X**_0_* = ***X-rowMeans(X)***. We then use ICA (‘*ica*’ function, ‘*icafast*’ library v1.0.2) or PCA (‘*prcomp*’ base R function) to decompose the data into a component space of dimension ***c*** such that ***X_0_ = G S^T^***, with ***G*** (***m × c***) the gene loadings and ***S*** (***n × c***) the sample scores. Columns of ***S*** are interpolated on with respect to time (and other potential variables of interest, *e.g.* batch), forming a new matrix ***T*** (***l × c***) of ***l*** new time points in component space. The full interpolated expression matrix ***Y***(***m × l***) is then reconstructed by multiplying the gene loadings matrix by the transposed ***T*** and by adding the gene centers ***Y = G T^T^*** + ***rowMeans(X)***.

To interpolate the components, we fit Generalized Additive Models (GAMs) to handle non-linear dynamics through splines with the ‘*gam*’ function in the ‘*mgcv*’ package (v 1.8.31) using a single model formula for all components selected by Cross-Validation (CV) as following: CV training sets are built with 80% of samples, with proportional representation of any covariate group (e.g. batch). The model is evaluated using the average relative error, mean squared error (MSE), and average root MSE^51^. We compared GAMs fitted with different splines (cubic, thin plate, duchon), and chose the model with minimal CV and prediction errors. Automatic spline parameter estimation from ‘*gam*’ function was used, unless the model was clearly performing poorly with automatic parameter estimation (overfitting, predictions not matching the component dynamics), in which case we performed further CV on reasonable spline parameter spaces to tweak the model (defining a number of knots). We further verified that RAPToR age estimates match chronological age of the original reference data and of independent time-series when staged on the interpolated reference, using the R^2^ of linear models (Sup. Note 1).

The number of components to fit was selected by setting a cutoff on cumulative explained variance (e.g. 99%). The cutoff was adjusted according to the number of components with intelligible dynamics with respect to time (Sup. Note 1, Sup. Fig. 8). Interpolation (and subsequent staging) is robust to variation in the number of components used (Sup. Note 1).

We implemented reference interpolation with the ‘*ge_im*’ function in the ‘RAPToR’ package. Model formulas and parameters for building all the references used in this study are displayed in Sup. Table 1.

### Age estimation

To perform **<u>a</u>**ge **<u>e</u>**stimation, we implemented the ‘*ae*’ function that takes the gene expression matrix to stage (genes as rows, samples as columns), the reference matrix (genes as rows and time points as columns), and the reference times (time values associated with the columns of the reference matrix) as inputs. The ‘*ae*’ function then finds common genes between sample and reference and computes the Spearman correlation between each sample and each reference time point. The age estimate for each sample is simply the reference time point with the highest correlation.

When an age estimate lands within 5% of the reference’s edges, we implemented a warning suggesting to stage the samples on another appropriate reference if possible.

To compute confidence intervals on age estimates, staging is repeated on bootstrap gene samples of default size of one third of the total. Unless stated otherwise, the number of bootstraps is 30. A confidence interval is given by the median absolute deviation (MAD) of bootstrap estimates (***est_boot_***) from the global estimate (***est***), and the resolution of the interpolation (***res***, time interval between 2 points of the interpolated reference):

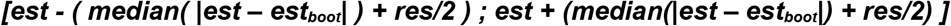

### Staging using a prior probability

We implemented the possibility of providing a prior probability in the form of parameters for a gaussian distribution per sample (mean, sd) which must be given in the time scale of the reference. A gaussian density function over the reference time is defined per sample from these parameters. During staging, all correlation peaks of the profile are determined and ranked by averaging their scaled correlation score (height of the peak in the correlation profile scaled to ***[0, 1]***) and prior score (value of the gaussian density function scaled to ***[0, 1],*** at the peak time point). The first peak of the ranking is then kept as the estimate. Since the ranking is determined by averaging normalized priors and correlation scores, changing the prior standard deviation parameter results in scaling the importance of the prior with respect to the correlation information.

No priors were used for staging unless explicitly stated.

## Evaluating RAPToR performance

### Staging C. elegans larval development

We built the reference from a time-series of WT larval development at 20°C sampled at 26 time points from L1 feeding to 48 hours^20^ (see Sup. Table 1), we set the number of interpolated time points to 500.

Staged samples are WT *C. elegans* collected during mid to late larval development at 25°C from 22 to 37 hours after L1 feeding^26^. Only samples aged below 32 hours (corresponding to about 48 hours at 20°C) were staged, to stay within the reference boundaries.

### Staging D. melanogaster embryonic development

We staged a *Drosophila* developmental time-series^27^ on an interpolated reference from another embryo developmental time-series^25^ (Dme_embryo reference of the drosoRef package, see Sup. Table 1). Samples were discarded when the 99^th^ percentile of the distribution of their Spearman correlation coefficients with others samples fell below 0.6, leaving 90 samples to stage. The number of interpolated time points in the reference was set to 500.

We compared our rankings with the BLIND^19^ rankings provided in the supplementary data^27^ (restricting to 77 samples as the authors used a more stringent quality cutoff).

To test if our age estimates better capture physiological development than chronological age, we fit identical linear models using the ‘*lmFit*’ function of ‘*limma*’ with either chronological age or RAPToR estimates as the predictor. Age is modeled using a natural cubic spline with 2 to 8 degrees of freedom (built with the *ns* function of the *splines* package). For each gene, we use R^2^ to compare the goodness of fit of the models with chronological age or RAPToR age estimates.

### Staging D. rerio embryonic development

We used the interpolated reference we built from embryo and larval development data^23^ (Dre_emb_larv reference of the zebraRef package, see Sup. Table 1) to stage a zebrafish time course of embryonic development from fertilization to 72 hours post-fertilization^27^. Samples were discarded when the 99^th^ percentile of the distribution of their Spearman correlation coefficients with others samples fell below 0.6, leaving 93 samples to stage. The number of interpolated time points in the reference was set to 1 000.

We then used the same reference, increasing the interpolation resolution between 0 and 15h to 800 time points (resulting in a reference time density of around 1 time point per minute instead of the previous 1 time point per hour) to stage an additional dense embryonic time-series of 180 zebrafish embryos around gastrula^28^. We compare RAPToR staging to rankings (Sup. Fig. 2a) previously determined^28^ as following: the 10 youngest and oldest embryos (determined through the morphological criterion of epiboly coverage) are used to select the genes with the largest decrease in expression from start to end of the time course. The average expression of these genes then determines the ranking. To show the benefit of reference interpolation, we also staged the embryos on the non-interpolated reference time-series (Sup. Fig. 2c-d).

### Staging M. musculus embryonic development

We used the interpolated reference we built from mouse embryonic development time-series data^24^ (Mmu_embryo reference of the mouseRef package, see Sup. Table 1) to stage an independent mouse somite-staged developmental time course29. The number of interpolated time points was set to 500. We compare RAPToR staging with the provided embryos somite number as no chronological age is given^29^.

### Staging M. musculus first-molar embryonic development

First and second data replicates for mouse first molar embryonic development are from Pantalacci et al.^37^, and Sémon et al.^38^ respectively. Reads from both replicates were processed together, trimmed with *trimmomatic*^52^ (v0.39) to remove adapters, and mapped using *salmon*^53^ (v0.14.1) and the Ensembl 98 version of the mouse transcriptome to obtain TPM values.

Genes with a median expression of log(TPM+1) < 0.5 across all samples were filtered out, leaving 15362 genes. A reference was built from both replicates of the lower jaw samples (see Sup. Table 1) and used to stage all 32 samples.

### Estimating developmental speed factors and resolution increase factors

Developmental speed factors and R^2^ between chronological and estimated age of samples are estimated with linear models.

We call ‘resolution increase factor’ the factor between sampling frequencies of a reference prior to interpolation and of a successfully staged independent time-series.

*C. elegans* larval development is sampled every 2 hours at 20°C (0.5/h) in the reference^20^ and every hour at 25°C (1/h, 1.5 development speed factor) in the staged time-series^26^ resulting in a resolution increase factor ***rf = (1.5 * 1)/0.5 = 3***.

*Drosophila* embryo development is sampled every 2 hours (0.5/h) in the reference^25^ and every 15 min (4/h) in the staged time-series^27^, resulting in a resolution increase factor ***rf = 4/0.5 = 8***.

Mouse embryo development is sampled every 1.5 days (0.66/day) in the reference^24^ and somite-staged in the target time-series^29^. Since the first 30 somites of *M. musculus* grow in ~2.5 days^54^, the somite-staged times series has a resolution of 12 time points per day (12/day) determining a resolution increase factor ***rf = 12/0.66 = 18.2***.

Zebrafish embryo development is sampled every hour (1/h) in the reference^23^ and at a rate equivalent to 47 per hour (47/h) in the staged samples^28^ (180 samples are roughly evenly staged between 5.7 and 9.5 hours post-fertilization: ***180 / (9.5 - 5.7)*** = 47/h), resulting in a resolution increase factor ***rf = 47***.

### Building aging references

To build RAPToR references capable of staging adults across independent studies, we select genes with monotonous expression along chronological age. Monotonous genes are defined as those with Spearman correlation with chronological age above a threshold given for each dataset. A threshold of √(0.33) selects genes where approximately a third of expression variance is explained by aging progression. When less than 200 genes were kept by the filter, we used a more lenient threshold of √(0.25). We then interpolate as described above, using only the first component (which is monotonous given the gene selection).

### Staging C. elegans aging

We built a *C. elegans* aging reference with an unpublished time-series (GSE93826) using monotonous genes, defined as those with spearman correlation with chronological age above √(0.33) (see Sup. Table 1). The number of interpolated time points was set to 500.

We then staged an RNASeq bulk time-series^33^, and a single-worm microarray profiling^34^. Single-worm behavior was provided by the authors^34^.

### Staging D. melanogaster aging

Pletcher et al.^35^ profiled aging *Drosophila* in food *ad libitum* and caloric restriction conditions. We built an aging reference with the ad libitum food time-series using monotonous genes, defined as those with spearman correlation with chronological age above √(0.33) (see Sup. Table 1). The number of interpolated time points was set to 500. We then staged all flies (control and caloric restriction).

### Staging human aging from brain tissue

We removed outliers from Chen et al.^36^ expression data when the 99^th^ percentile of the distribution of their Spearman correlation coefficients with other samples fell below median+2 standard deviations, and when the reported RNA integrity score was below 7. The remaining 360 samples were split per tissue (BA47 or BA11), and genes with spearman correlation with chronological age above √(0.25) were defined as monotonous. For each tissue, half of the samples were randomly selected to build a RAPToR reference with monotonous genes (see Sup. Table 1), on which all samples were then staged. Samples were also staged on the reference built from the other tissue (Sup. Fig. 10).

### Staging H. Sapiens single-cell embryo development

We filtered Petroupoulos et al.^39^ single-cell counts to remove genes with a median expression of log(TPM+1) < 0.5 across all samples, leaving 8482 genes. 20% of cells were randomly sampled from each of the 5 time points (305 cells) to build an interpolated reference (see Sup. Table 1), on which all 1529 cells were then staged (only non-reference cells are shown in Fig. 2l). The cell pseudo-times are from the original study^39^, provided as sample metadata in the ArrayExpress entry.

### Probing robustness of reference interpolation

Robustness of reference interpolation to the choice of dimensionality reduction method and number of components was evaluated using either the *C. elegans* time-series by Kim et al.^20^ (as above), or the one by Meeuse et al.^21^ as references.

Robustness was evaluated computing Sum Squared (SSQ) of gene expression prediction error by reference models using PCA or ICA and 2 to 16 components with the Kim et al. time-series, and 2 to 20 with the Meeuse et al. one. The model formula was fixed to the one defined in Sup. Table 1. The SSQ prediction error is defined as ***SSQerror = Σ((X_(n x m)_ − X_pred_)^2^) / (n * m),*** with ***n*** samples, ***m*** genes.

For 6 conditions – ICA/PCA, each at 3 different numbers of components – we staged the reference samples as well as an independent *C. elegans* time-series^26^ on the interpolated reference (only samples within reference boundaries were staged on the Kim et al. reference). We evaluated models built from 4, 9, and 14 PCA or ICA components for the Kim et al. reference and models built from 10, 20 and 25 PCA or ICA components for the Meeuse et al. reference.

We then reported the R^2^ value of a linear fit of RAPToR estimates by the chronological age of the samples in each condition (Sup. Table 2), as well as the correlation score between the samples and the interpolated reference at the estimate (Sup. Fig. 7).

### Estimating the impact of gene set size on staging

The impact of the gene set size on staging was evaluated by staging the *C. elegans* larval time-series by Hendriks et al.^26^ on the reference built from the Kim et al.^20^ samples, as above.

We staged the samples using 50 random gene sets of sizes 16 000, 12 000, 8 000, 4 000, 2 000, and 1 000. The resulting estimates were used to compute confidence intervals for varying bootstrap set sizes. We reported the median absolute deviation of estimates to the full gene set estimate plus interpolation resolution (i.e. the size of half the confidence interval).

The same approach was repeated for smaller gene set sizes of 2 000, 1 000, 500, and 250, this time staging the samples with and without priors (defined as 1.5 times the chronological age of the samples to account for the developmental speed difference with the reference; prior standard deviation was set to 10).

### Tissue-specific staging and quantification of soma-germline heterochrony

208 recombinant inbred lines (RILs) from a cross between N2 (Bristol) and CB4856 (Hawaii) strains of *C. elegans* were genotyped at 1,455 SNP markers, and collected as young adult hermaphrodites (originally intended as one time point) for profiling by microarray with one sample per RIL^11^.

Microarray intensities were first normalized within arrays with LOESS using the ‘*normalizeWithinArrays*’ function of the ‘*limma*’ library. Arrays corresponding to pooled mixed stage controls were then discarded. Samples were discarded when the 99^th^ percentile of the distribution of their Spearman correlation coefficients with others samples fell below 0.95, leaving 193 samples for analysis.

The reference used to stage the samples is the “Cel_larv_YA” reference^21^ of the wormRef package (see Sup. Table 1). The number of interpolated time points in the reference was set to 1000.

Samples were first staged using the entire available gene set to obtain the global estimates, then with somatic and germline specific gene sets to obtain the corresponding tissue-specific estimates: the somatic gene set corresponds to the oscillatory genes denoted “osc” in Hendriks et al.^26^. The “germline” gene set corresponds to the union of “germline_intrinsic”, “spermatogenesis_enriched”, and “oogenesis_enriched” gene sets defined in Reinke et al.^26^. Estimating somatic age, required the use of the global estimate as prior (due to gene expression oscillations generating multiple correlation peaks), with the prior standard deviation set to 10 for all samples. Germline age estimates required no prior.

To compare expression dynamics between reference and RILs, we kept the overlapping genes between the non-interpolated reference and the samples, quantile-normalized both datasets together, and performed an ICA (‘*ica*’ function of ‘*icafast’)* extracting 46 components, explaining 95% of the variance in the joined data. For components 2-8, capturing developmental signal (IC1 captured batch effect, IC9 captured genetic variation), we defined genes with an absolute value of loadings above 1.96 as contributing to the component, and tested for their enrichment of soma, oogenesis and spermatogenesis genes with a two-sided hypergeometric test, whose p-values were adjusted across all tests with the Benjamini-Holm method.

To test heterochrony between RILs and the reference, we fit splines on the reference samples in IC2-IC8 (using the same model as the RAPToR reference, see Sup. Table 1) and computed Root Mean Square Error (RMSE) between the fit and RILs for each component. This was done for global, soma and germline age estimates of RILs (Fig. 3d), and for shifted values of global age estimates (−5h to +5h, Sup. Fig. 13c).

To test the existence of heterochrony among the RILs, we fit identical models on the RIL expression data using the ‘*lmFit*’ function in *limma* with global, soma, or germline age values as predictors. We used natural cubic splines (‘*ns*’ function in the ‘*splines*’ library) on the age with 4, 6, or 8 degrees of freedom. Choice between models (at equal spline degrees of freedom) was done per gene based on highest R^2^ value.

#### Quantitative Trait Loci (QTL) analysis on soma-germline heterochrony

The multivariate QTL analysis on soma-germline heterochrony among RILs defined as (soma age) - (germline age) was performed by Random Forest (RF) regression^55^ with or without batch as a covariate. Each RIL was genotyped at 1455 SNP markers^11^. Redundant markers were filtered out from the selected 193 RILs, missing values for the remaining 1105 markers are imputed with the ‘*rfImpute*’ function and random forest regression was fit with 5000 trees using the ‘*randomForest*’ function; both functions are from the ‘randomForest’ package (v4.6.14). The RF Selection Frequency (RFSF) was used as importance measure, adjusted for selection bias^55^ which was estimated by fitting 500 forests of 10 trees to gaussian noise.

We estimated the null probability distribution of RFSF through 100 trait permutations, calculated empirical p– values and adjusted them for FDR.

### Cross-species staging

#### Staging non-model Drosophila on D. melanogaster

We used the interpolated reference we built from the *D. melanogaster* embryo development^25^ (Dme_embryo reference of the drosoRef package, see Sup. Table 1) to stage time courses of development of 6 *Drosophila* species^41^: *Drosophila melanogaster, simulans, ananassae, pseudoobscura, permisilis* and *virilis* profiled by microarrays. We used orthologs provided by the authors^41^. The number of interpolated time points in the reference was set to 500.

Developmental speed difference from *D. melanogaster* was determined with a linear model without intercept predicting RAPToR estimates with the chronological age of samples, with species as covariate and including interaction. Comparison with the original scaling factors^41^ is shown in Sup. Table 3.

To determine whether RAPToR estimates or the linearly-scaled age from the study^41^ is the better development indicator, we fit identical linear models on gene expression (*lmFit* function of *limma*) with either values as the predictor, and species as covariate. Age is modeled using a natural cubic spline with 2 to 8 degrees of freedom (*ns* function of *splines*). For each gene, we use R^2^ to compare the goodness of fit of either model (Sup. Fig. 16). No interaction between age and species coefficients was considered as temporal scaling of development between species is already applied.

We evaluated the effect of species distance on staging through the maximal correlation coefficient between the samples and the reference (i.e. at their age estimate).

#### Staging C. elegans on Drosophila

We staged a *C. elegans* embryo time-series^27^ on the interpolated reference we built from the *D. melanogaster* embryo development time-series^25^ (“Dme_embryo” reference, drosoRef package). First, poor quality *C. elegans* samples were discarded when the 99^th^ percentile of the distribution of their Spearman correlation coefficients with others samples fell below 0.67. Additionally, a sample (GSM1487346, or “sample_0029”) was also excluded as it clearly appeared as an outlier on multiple ICA components (Sup. Fig. 19). 4 samples (GSM1487318, GSM1487319, GSM1487320, GSM1487321, or “sample_0001” through “_0004”) were further removed due to erroneous chronological age (Sup. Fig. 19), leaving 127 samples.

We then performed the staging using a restricted fly-worm ortholog set^45^. We also did staging on a second reference interpolated as above but using the first 2 instead of 8 components. For both interpolated references, the number of interpolated time points was set to 500.

Further analysis is restricted to the overlapping set of orthologs between worm and fly datasets (3194 genes). We ranked genes by Spearman correlation between the *C. elegans* embryo time-series and their matching timepoints in the second *D. melanogaster* reference. We then selected the 10% genes with highest correlation (319 genes) and staged the *C. elegans* samples once more on the second *D. melanogaster* reference, evaluating staging performance with Spearman correlation and the R^2^ of a linear model between chronological age and estimated age.

Hierarchical clustering the top 10% genes in the original *D. melanogaster* reference data^25^ *(‘hclust*’ function on the euclidean distance matrix of gene-centered log(TPM+1)), resulted in 3 clusters with over 20 genes. We then evaluated gene ontology enrichment in each cluster with gProfiler^56^ using the 3194 overlapping set of worm-fly orthologs as background (Sup. Table 4, 5, 6).

#### Staging M. musculus on H. sapiens

We filtered Deng et al.^44^ mouse early-embryo single-cell counts to remove genes with a median expression of log(TPM+1) < 0.5 across all samples, leaving 6506 genes. We then staged cells using all available mouse-human orthologs from ensembl, on the interpolated reference we built from *H. sapiens* embryo development single-cells^39^ (see *Staging H. Sapiens single-cell embryo development* and Sup. Table 1). We compared age estimates with chronological age, as well as with pseudotime rankings similar to Petropoulos et al.^39^ (Sup. Fig. 18): we fit a principal curve (*principal_curve* function of *princurve* library) on the first 3 components of a PCA on the 1000 most variable genes.

As done above for worm-fly staging, we then restricted further analysis to the overlapping set of orthologs between mouse and human datasets (6057 genes), ranked genes to select the 10% genes with highest correlation between both species (509 genes), and staged the *M. musculus* single-cells once more on the human reference, evaluating staging performance with Spearman correlation and the R^2^ of a linear model between chronological age and estimated age.

Hierarchical clustering of the top 10% genes in the original *H. sapiens* reference data (expression matrix aggregated per sampled time point, gene-centered log(TPM+1)), resulted in 5 clusters with over 20 genes. We then evaluated gene ontology enrichment in each cluster with gProfiler^56^ using the 6057 overlapping set of mouse-human orthologs as background (Sup. Tables 7-11).

#### Staging H. sapiens on B. taurus

We filtered an RNAseq cow early-embryo time-series^43^ to keep genes with median log(TPM+1) expression >0, and built a reference with half of the samples. We similarly kept genes of a microarray human embryo time-series^42^ with log(X +1) expression > 2 and built a reference with half the samples (see Sup. Table 1). As only morphological stages (e.g. 2-cell, 4-cell) and not timings were given in both datasets, we used timings from the literature^57^. The number of interpolated time points for both references was set to 100. We then staged all cow samples on the human reference and vice versa, using all available human-cow orthologs from ensembl (Sup. Fig. 17).

### Exploiting RAPToR age estimates

#### Drug dose response on developmental delay in *C. elegans*

Expression profiles of young *C. elegans* adults exposed to drugs^46^ were staged on the “Cel_larv_YA” reference^21^ from the wormRef package (Sup. Table 1), with 500 interpolated time points in the reference. We estimated global, soma-specific, and germline-specific ages (see Tissue-specific staging). For each age type, we then subtracted the age of the control sample within each replicate of each drug assay to compute the developmental difference by treatment group. We fit a linear model with drug, dose, and interaction on the age differences to assess the significance of the effects.

#### Increasing statistical power in differential expression analyses

WT and *pash-1ts C. elegans* samples^47^ were staged on the “Cel_YA_2” reference^22^ from the wormRef package (Sup. Table 1), with 500 interpolated time points in the reference. The second replicate of the first wild-type time point (wt_h0.2) was omitted from further analysis due to its extreme developmental displacement and lack of comparable mutant sample.

We fit identical linear models with the ‘*lmFit*’ function in the ‘*limma*’ library to test for differential expression, including either chronological or estimated age modeled with a natural cubic spline (‘*ns*’ function in ‘*splines’,* df = 2), strain and their interaction.

Effect of strain and development was then assessed by considering the significance of appropriate model coefficients (interaction and strain coefficients for strain effect, spline and interaction coefficients for development effect), with the ‘*topTable*’ function in the ‘*limma*’ library. Differential expression was considered significant at 0.05 Benjamini-Hochberg False Discovery Rate (FDR).

To test the effect of similar random age differences from chronological age, we generated 100 “random age” sets by sampling age differences from the distribution of (chronological age) - (estimated age) values, estimated with the ‘*density*’ function in R. Sampled age differences were then added to the chronological age, and the same model and analysis as above was applied. The goodness of fit per gene is assessed using R^2^.

#### Quantifying developmentally-driven gene expression changes

Given any two groups of expression profiling samples ‘A’ and ‘B’, we first stage them, then fit a linear model per gene on log_2_(TPM+1) (or log2(Intensity+1) for microarray expression data) to compute the observed log_2_-fold changes of ‘A’ vs. ‘B’ samples. Then we fit the same model on reference profiles at matching time points to compute log2-fold changes expected from development only (Sup. Fig. 23) and we use squared Pearson correlation between observed and expected logFCs to quantify the variance explained by development in the observed logFC.

Control and post-dauer *C. elegans* samples^48^ were germline-staged (see *Tissue-specific staging*) on the “Cel_larv_YA” reference^21^, and on the “Cel_YA_2” reference^22^ of the wormRef package for confirmation, as they landed near the edges of the first reference. The number of interpolated time points in the Cel_larv_YA and Cel_YA_2 references were set to 1000 and 500 respectively. Using the method described above, we quantified the differential expression explained only by difference in developmental stages between the control and post-dauer samples.

We could not compare our results to the original results as we were unable to exactly reproduce the distribution of DE and p-values of the original t-test based analysis. We therefore recalculated DE gene expression using linear models (function ‘*lmFit* in ‘*limma*’ library in R).

#### Recovering direct perturbation effects using reference data

WT and *xrn-2* time-series of *C. elegans* late larval development^49^ were staged on the “Cel_larv_YA” reference^21^ from the wormRef package (Sup. Table 1), with 500 interpolated time points. We restricted further analysis to the genes with both ≥5 raw counts for at least one sample, and overlapping with the reference gene set (17656 genes).

##### Defining the differential expression gold standard

To establish the gold standard of DE genes, we selected time points 8 to 10 of *xrn-2* and WT, as they had the best (estimated) developmental match. We then calculate differential expression fitting a generalized linear model (GLM) on raw counts using the *glmFit* function of *egdeR* (v3.28.1), including only the strain variable (model 1), and considered genes DE with Bonferroni-Holm adjusted p-values < 0.05 of a likelihood ratio test *(glmLRT* function of ‘*edgeR’)* on the strain coefficient.

##### Evaluating gold-standard gene detection decrease with age gap

To test how increasing mismatch in developmental time between *xrn-2* and WT impacts DE analysis we apply the same GLM used for the gold standard (model 1) to calculate differential expression between the mutant and WT samples shifted by −1, −2, −3, −5, and −7 time points and we estimated expression changes explained by development as detailed above (*Quantifying developmentally driven gene expression changes*). We then evaluated how well model 1 p-values detect gold standard DE genes at increasing age gaps by Precision-Recall Curves (PRC) and area under PRC using the ‘*prediction*’ function of the ‘*ROCR*’ package (v1.0.11).

##### Correcting expression changes from development

To accurately account for developmental changes we combine the samples of interest with the interpolated reference.

For each set of samples (including WT and mutant samples), we define the window of reference to include as the range of age estimates widened by a 1 hour margin on either side. For example, in the ‘WT-1’ set, the youngest sample (WT_05h) is 51.7h old, and the oldest (xrn.2xe31_09h) 58.3h old. Thus, we include the interpolated reference from 50.7h to 59.3h of development.

We transform the interpolated reference data to artificial counts assuming a fixed library size of 25*10^6 counts per sample and a fixed number of reads “per gene length” defined by the median of available gene lengths:

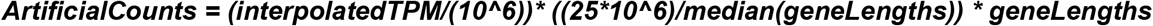

The artificial count matrix is then joined to the sample count matrix, and a GLM is fit (‘*glmFit*’ in ‘*edgeR”*), including batch (between reference and sample data), the variable of interest (strain) where reference data is grouped together with the control, and developmental time modeled with splines (‘*ns*’ function in ‘*splines*’). To select the optimal spline degree of freedom for each window, we minimized the residual sum of squares of a linear model fit on the reference window only (Sup. Fig. 24h). Only model coefficients of the variable of interest (strain logFCs) are considered.

We first evaluated how well strain logFCs detects DE genes from the gold standard using PRC and AUPRC *(‘prediction*’ function in ‘*ROCR*’). We then defined an Age-Corrected Classifier (ACC) as the weighted mean of the model 1 p-value and strain logFC of the model including the reference:

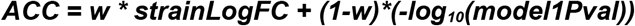

with ***w***, the weight ratio of either classifier. We defined the optimal ***w*** as the value for which the area under the precision recall curve (AUPRC) is maximal, and estimated it for each set of WT shifts. At optimal ***w***, we then reported the AUPRC of our age-corrected classifier and compared it to the standard model.

As the optimal ***w*** cannot usually be estimated in this way, we explored the relationship between optimal ***w*** and the correlation between observed and expected logFC (as defined in *Quantifying developmentally driven gene expression changes)* calculated for a larger amount of WT 3-sample sets (Sup. Table 13).

## Acknowledgements

We are grateful to Sarah E. Hall, Marie Sémon, and Sophie Pantalacci for providing data from their profiling experiments. We are also grateful to Gael Yvert, Daniel Jost, Marie Sémon, Sophie Pantalacci, and Ben Lehner for their critical reading of the manuscript.

## Author Contributions

MF and RB conceived the method, RB developed the computational framework and performed the analyses, MF and RB wrote the manuscript.

