## Supplementary material for "Real age prediction from the transcriptome with RAPToR": Suplementary Notes and Figures

### Supplementary note 1

#### Computational Requirements

RAPToR is an R package. We have tested and confirmed the package works in R 3.6.3 and 4.1, under Windows 7 & 10, Ubuntu 18.04 & 20.04, and macOS (10.14). The package installs under 20s once dependencies are met. Standard datasets can easily run with 4Gb of RAM and 2 CPU cores. For reference, the dataset used for demo in the main documentation vignette of the package (43 samples by ~19500 genes, GSE80157 [1]) can be both downloaded and staged with RAPToR in under 30 seconds, using less than 2Gb of RAM.

#### Reference interpolation

For reference interpolation, we take advantage of the redundancy of gene expression. We use either Principal or Independent Component Analysis (PCA, ICA) to decompose the expression matrix into components (sample scores, or eigengenes) that summarize the gene expression dynamics, and gene loadings that are gene contributions to each dynamic [2,3]. We interpolate the components with respect to time (Fig 1e), and reconstruct the full interpolated gene expression by the matrix product of interpolated components and the gene loadings (Fig 1f, methods). In this way, we simplify model building and validation across thousands of genes to a few components. We select a number of components by a threshold on cumulative variance explained (see methods), keeping components with ‘intelligible’ dynamics which we defined as those with a spline fit explaining > 0.5 of the deviance (Sup. Fig. 8). However, staging results are robust to the variation number of components used (Sup. Fig. 7a, 7b, see below).

In order to validate an interpolated reference for staging with RAPToR, we first stage the original reference data on its interpolated version. We fit a linear model predicting RAPToR age estimates with chronological age expecting a near-perfect match, with adjusted $R^2 > 0.99$, and non-significant intercepts. This was the case for all references presented throughout the study, aside from “Cel_YA_2” which is built from old data[4] (2004) and only had an $R^2$ of 0.901. Then, when possible we stage independent time course experiments and expect very good fits of linear models ($R^2 \geq 0.9$, as shown in Fig. 2). We note that increased variability is expected from single-organism profiling data (such as Fig. 2d and Fig. 2h) as well as for aging time-series (such as Fig. 2j and Sup. Fig. 10), due to their inherent biological heterogeneity.

Interpolation errors or bias are more frequent at the edges of a time series (e.g. splines are known to behave erratically at edges). In addition, when the real age of a sample is outside the reference range, it will likely match a time point close to the edge, making estimates close to the edges less reliable. Therefore, when age estimates are found near the edges, we suggest using another overlapping reference to confirm the estimates.

#### Evaluating RAPToR performance

##### *Reference interpolation effectively increases estimates temporal resolution*

RAPToR uses correlation with a reference to estimate age. If we were to use a non-interpolated reference (ie. the expression matrix in Fig. 1d), this would only allow estimates at the sampled time points of the reference. Reference interpolation enables age estimates between the original reference time points.

We successfully staged worm[5], fly[6], and mouse[7] time-series (Fig. 2a, 2d, 2c) with 3, 8, and over 18 times the temporal resolution of their respective original references (see methods). This clearly shows that interpolation allowed us to accurately reconstruct gene expression dynamics of the reference time-series at an increased temporal resolution and consequently improve the accuracy of the age estimates.

To further test this, we staged another zebrafish time-course consisting of 180 embryos spread in a very short developmental window around gastrulation[8]. Despite having over 40 times the estimated temporal resolution of the reference before interpolation, the data is accurately staged by RAPToR, not only matching the established ranking[8] ($\rho$=0.99, Sup. Fig. 2a) and expression dynamics (Sup. Fig. 2b), but giving absolute times that are comparable to any estimate obtained with the same reference.

We additionally staged the same zebrafish samples on the non-interpolated reference data to explicitly show the gain from interpolation. Samples have the highest correlation with the expected reference time points, still matching the aforementioned ranks ($\rho$=0.94, Sup. Fig. 2c) and our previous estimates ($\rho$=0.94, Sup. Fig. 2d), but the staging is imprecise due to the reference data only having one expression profile per hour of development.

##### *Effect of gene set size and data quality on age estimates and bootstrap intervals*

We tested RAPToR robustness to changes in the gene set size by staging an independent time series[5] of C. elegans larval development on the reference[9] we built (as in Fig. 2a). With random gene sets of 4000 genes, estimates have a median absolute deviation under 10 minutes from full gene set estimates (18718 genes);

with sets of 1000, under 20 minutes (Sup. Fig. 3a, 3b). With smaller sets (<1000), estimates are unreliable likely because repeated expression patterns – such as oscillations or pulses – create multiple correlation peaks with the reference (Sup. Fig. 3c). For example, the repeated molts of *C. elegans* larval development generate oscillations in the correlation profiles of staged samples (Sup. Fig. 4). To address this, we implemented the possibility to include a prior for estimates (see methods). With the prior, no estimates get misplaced to other correlation peaks, even for sets of a few hundred genes (Sup. Fig. 3c, 3d). We note that priors must be given in reference time (though approximate timings are enough, within a few hours), which is why we provide correspondence between developmental stages and chronological time in the references through RAPToR (*plot_refs* function).

Potential sources of uncertainty for staging can be technical (profiling quality, number of genes available), or biological (developmental spread of individuals within a bulk sample, heterochrony between tissues). We explored how these factors could influence the confidence intervals (CI) of RAPToR estimates using the *C. elegans* time-series[5] and reference[9] mentioned above. We find that noise in reference data gets largely filtered out by the PCA, and thus does not impact the CI size (Sup. Fig. 6a). Staged samples of poor quality have wider CIs, but estimates stay strikingly accurate with intense noise (Sup. Fig. 6c). Indeed, even when noisy expression profiles have low correlation scores with the original data ($\rho<0.8$, Sup. Fig. 6d), staging accuracy is not diminished ($R^2=0.99$, Sup. Fig. 6c). As discussed above, with less information (fewer genes) the reliability of staging decreases, which leads to increasing CIs. To simulate developmental spread within a sample, we averaged the expression values of 2, 3, 5, and 7 consecutive time points in the staged time-series; similarly, we simulated heterochrony by randomly selecting one of the expression values between 2, 3, 5, and 7 consecutive time points (i.e. mixing). As expected, CI size increases in both scenarios (Sup. Fig. 6e, 6f).

We conclude our bootstrapping approach generates confidence intervals that reflect the multiple technical or biological sources of variability in the data.

Confidence intervals of RAPToR estimates for *C. elegans* samples[5], built from bootstrap estimates with 6239 genes (see methods), are extremely small – between 5 and 20 minutes (Sup. Fig. 5a). As expected, staging zebrafish embryos[6] on a reference[10] with lower gene overlap (8662 genes) results in larger confidence intervals built from bootstrap estimates with 2887 genes – on average, slightly over 2h across the full time series, and under 50 minutes for samples before 30h of development (Sup. Fig. 5b).

Of note, the combination of interpolation accuracy and estimate precision for small gene sets means that even time-series transcriptomic data with low sampling rate or gene coverage can be exploited to build interpolated references.

##### *Effect of interpolation parameters on age estimates*

We tested whether reference-building is robust to parameter changes. As expected, selecting more components leads to a decrease in prediction error of the model (Sup. Fig. 7a, 7b), and a slight increase in the correlation between staged samples and reference at estimate (Sup. Fig. 7c, 7d). However, the age estimates and bootstrap intervals were mostly unperturbed by the changes in number of components and were also robust to choosing PCA or ICA for interpolation (Sup. Table 2, Sup. Fig. 6b).

Scenarios such as aging and cross-species staging can require building references with few, or even a single component. In this case, expression dynamics are still accurately interpolated when they can be reconstructed from linear combinations of the selected components. For example, using a single monotonous component to build a reference will result in genes with complex expression dynamics being poorly modeled, while those with monotonous dynamics will have accurate fits.

##### *Within-study aging expression dynamics allow robust RAPToR staging*

While there is a known increase of gene expression noise with aging[11], the heterogeneity of aging also greatly depends on environmental factors. Indeed, we find that even though "standard" (whole-transcriptome) RAPToR references poorly stage independent experiments, they accurately stage independent samples from the same study. We show this in *C. elegans* ($R^2$>0.99, Sup. Fig. 9a) and even in dissected tissues of wild-type and transgenic mice[12] ($R^2$>0.99, Sup. Fig. 9b). Together with the fact that RAPToR estimates stay accurate in very noisy expression data (see above, Sup. Fig. 6c), this suggests that the reason for poor staging performance across studies is likely due to heterogeneous environmental conditions across studies.

##### *RAPToR stages human adults from dissected brain tissue*

RAPToR aging references built with monotonous genes allow us to infer the age of adults accurately despite heterogeneous environments and genetic backgrounds. In humans – where neither of these factors is controlled – we could accurately stage individuals from the transcriptome of dissected tissues of two neighboring brain regions[13]: Brodmann Area BA11 ($R^2$=0.74, Sup. Fig. 10a) and BA47 ($R^2$=0.68, Sup. Fig.

10d). We could even stage BA11 samples on a BA47 reference ($R^2$=0.61, Sup. Fig. 10b) and vice versa ($R^2$=0.71, Sup. Fig. 10e) with comparable accuracy. Importantly, BA11 sample age estimates on the BA11 reference finely match those acquired on the BA47 reference ($R^2$=0.93, Sup. Fig. 10c) and the same goes for BA47 samples ($R^2$=0.93, Sup. Fig. 10f). This suggests that RAPToR captures genuine variability between chronological and physiological age, since the same age is given to a sample with both references.

In summary, RAPToR can infer the age of adult individuals with heterogeneous environmental and genetic backgrounds from the transcriptome of dissected tissues, and do so reliably from a reference built with a similar tissue.

##### *Inferring developmental speed factors*

Beside the expected 1.5 fold increase in developmental speed due to temperature we observed in *C. elegans* (Fig. 2a), we also observe a difference between chronological and estimated times in the independent zebrafish developmental time series[6] we staged on the zebrafish reference[10] determining a developmental speed factor of 0.7. We suspect this speed factor is also due to a temperature difference with the reference, as growth speed scales with temperature also in zebrafish. While the reference data embryos developed at 28.5°C[10] , we were unable to confirm at which temperature the staged data embryos developed but we presume a lower one.

##### *Soma-germline heterochrony between C. elegans experiments*

To confirm the presence of soma-germline heterochrony between *C. elegans* Recombinant Inbred Lines[14] (RILs) and the reference[15] they were staged on, we compared the expression dynamics of both datasets (Sup. Fig. 13a). While the overall Root Mean Square Error (RMSE) between the RILs and the reference fit is minimal at the RIL global estimated age (Sup. Fig. 13c), that is not the case for single components. Indeed, reference dynamics match RILs better when shifting the RIL age estimates back for soma-enriched components and forward for germline-enriched components (Sup. Fig. 13d). Soma- and germline-specific age estimates using tissue-specific genes then further improve the match between the RILs and reference for dynamics of these respective tissues (Fig. 3g)

Soma-germline heterochrony between the reference and RILs only explains that the dynamics of the soma be shifted along germline age and vice versa. However, the clear noise increase we see in germline dynamics along soma age (Fig. 3c) and vice versa (Fig. 3f), also implies heterochrony among RILs as it shows the soma-germline age difference with the reference varies from sample to sample.

##### *Quantitative Trait Loci analysis on soma-germline heterochrony*

In our QTL analysis of soma-germline heterochrony in *C. elegans* Recombinant Inbred Lines[14], Random Forest prediction of the trait was poor and non-significantly correlated with the trait (r = 0.12, p = 0.09) and we found no significant hits out of the 1,455 markers, even at FDR of 0.5. Removing the batch covariate from the analysis results in even poorer predictions (r = 0.08, p = 0.2), suggesting uncontrolled environmental factors may be driving heterochrony.

##### *Staging samples on references of a different specie*

When we first staged a *C. elegans* embryo development time course[6] on the *Drosophila* reference[16], we noted two breaks in the age estimates, possibly due to heterochrony of developmental processes between the 2 species ($R^2$ = 0.938, Sup. Fig. 19a). Staging was notably improved after rebuilding the reference with only 2 components which have broad dynamics ($R^2$ = 0.958, Fig 4d), further suggesting that sharper expression dynamics (like pulses or oscillations) diminished staging performance.

To explore the reason behind successful cross-species staging, we analyzed the 319 genes (10%) with highest correlation between *C. elegans* and *D. melanogaster* during embryogenesis, as well as the 509 genes (10%) with highest correlation between *H. sapiens* and *M. musculus* early embryo development in single-cells (see methods), which suffice to stage the embryos well despite their small number ($R^2$ = 0.87 and $R^2$ = 0.84, Sup. Fig. 19b and 18c). We found the fly-worm gene set clusters into an ascending gene expression signature of muscle development (cluster 1, Sup. Fig. 19c, Sup. Table 4), and two decreasing signatures of cell proliferation split between DNA replication (cluster 2, Sup. Fig. 19c, Sup. Table 5) and splicing (cluster 3, Sup. Fig. 19c, Sup. Table 6) respectively. Similarly, the mouse-human gene set consists of clear ascending expression signatures of cell respiration, adhesion and secretion (clusters 1-3, Sup. Fig. 18d, Sup. Tables 7-9) and of a strong decreasing signature of cell proliferation (cluster 4, Sup. Fig. 18d, Sup. Table 10). Other translational-related processes are grouped without a clear trend (cluster 5, Sup. Fig. 18d, Sup. Table 11)

Expression dynamics match well between fly and worm embryogenesis (Sup. Fig. 19c), and between human and mouse early-embryo development, consistent with previous work showing widespread conservation of decrease in cell proliferation during embryogenesis[6] .

### Supplementary note 2

#### Exploiting inferred age in genome-wide expression studies

##### *Inferring the impact of environmental or genetic perturbations on development*

When staging C. elegans exposed to increasing concentrations of mefloquine, dichlorvos, and fenamiphos[17], we noted that beyond the germline developmental delay induced by all three drugs, dichlorvos also showed a significant and opposite effect of dose on somatic age. However, the scale of the effect is a fraction of the one observed on the germline (Sup. Fig. 20b).

##### *Exploiting developmental variation to increase power to detect differential expression*

After comparing chronological age and RAPToR age estimates as predictors for *C. elegans* control and *pash-1* mutants profiled at 4 time points of late development[18] and finding better model fits (Fig. 5c), we further tested whether random perturbations on age could induce a similar result. Thus, we generated age sets of similar deviation from chronological age than RAPToR estimates (see methods) and found that these consistently decreased model fits and DE gene detection, confirming that precise age estimates increase the power of the analysis (Sup. Fig. 21a-c, methods).

We also found a curious batch effect on development: mutants are systematically older than controls in the first two replicates while it was the opposite in the third (Sup. Fig. 21d).

#### Detecting and correcting expression changes caused by development using reference data

When samples are few and experimental groups have little or no overlap in development, the information available in the experiment is not sufficient to separate the effects of development from those of interest. To overcome this, we developed an approach using RAPToR interpolated references to quantify and correct for development in genome-wide expression data.

##### *Quantifying developmental expression changes*

To quantify the impact of development in DE analysis, we compare observed $log_2$-fold changes (observed logFC) between the two groups (i.e. mutant and wt) with changes expected purely from developmental differences between groups (expected logFC) which we estimate comparing age-matched interpolated reference profiles (Sup. Fig. 23). We quantify development impact using Pearson correlation between observed and expected logFCs (or its square). We use Transcripts Per Million (TPMs) to compute the logFCs, as they are more comparable across samples and datasets.

##### ***Development can completely confound DE analysis leading to erroneous conclusions***

Not accounting for confounding developmental variation in DE analysis can lead to erroneous conclusions. Comparing young adult *C. elegans* that developed through dauer state (post-dauer) to controls that did not, Hall et al. [19] conclude that post-dauer animals have reduced spermatogenesis from a down-regulation of spermatogenesis-associated genes and an up-regulation of oogenesis-associated genes in their DE analysis. However, this could simply be explained by post-dauer samples being older than controls as *C. elegans* naturally switch from sperm to egg production during development. Moreover the increased brood size in post-dauer worms described by the authors[19] would even suggest the opposite as sperm number limits brood size in *C. elegans*: post-dauer animals would have up- and not down-regulation of spermatogenesis genes.

To rigorously test if these expression changes are caused by development, we estimated the global and tissue-specific age of samples with our best-quality reference, and found post-dauer samples were systematically older. However, while germline age estimates were reliable, global and soma-specific staging put some samples at the edge of the reference, indicating development beyond reference bounds (Sup. Fig. 22a). Thus, we validated the age estimates with an older and lower-quality reference spanning a few hours further and found the same divide in global, soma, and germline age between groups (Sup. Fig. 22b).

RAPToR age estimates show the control samples are in late spermatogenesis, while the post-dauer samples are 5-10 hours older, fully switched to oogenesis (Fig 5c, Sup. Fig. 22a). A DE analysis between the two conditions does recapitulate reported divide between spermatogenesis and oogenesis genes (Sup. Fig. 22c). However, this is also fully recapitulated by the expected developmental changes ($r = 0.82$, Fig 5d, Sup. Fig. 22d). Correlation also stands with all genes ($r = 0.44$, Sup. Fig. 22e). Repeating the logFC comparison with the older reference yielded similar results (germline $r = 0.74$, all genes $r = 0.41$ Sup. Fig. 22f, 22g).

This supports our hypothesis that the expression changes between groups, particularly in germline genes, is due to a difference in development between the samples, rather than a direct effect of the post-dauer condition.

##### ***Recovering direct perturbation effects using reference data***

To recover the effect of a perturbation confounded by development we propose a model in which we include all reference expression profiles in a time window spanning the development of the selected samples (see methods) and model expression dynamics as a spline of developmental time, batch between reference and sample data, and the perturbation of interest. We first convert interpolated reference data from TPM to counts, assuming a fixed library size (see methods) to ensure compatibility with tools that require counts to calculate differential expression. Including reference data with artificially low dispersion invalidates models statistics such as p-values. We can however rely on the model coefficients (logFCs) estimated with the reference data to account for development. The perturbation (strain) logFC coefficients estimated by this reference integrated model are controlled for developmental changes shared by the samples and the reference (Sup. Fig. 24a).

To evaluate how effectively our approach recovers truly DE genes, we exploit time series data of *C. elegans* *xrn-2* and WT late larval development sampled every hour at 25°C[20] . We first define a gold standard of truly DE genes by comparing three mutant and three WT samples with the best developmental match. Next we evaluate the effect of increasing developmental difference on the DE analysis by comparing the same three mutant samples with three increasingly mismatching WT samples (Fig. 5d, methods). We shifted WT samples back by 1, 2, 3, 5 and 7 time points (corresponding roughly to 1, 2, 3, 5, and 7 hours of development at 25°C). Correlation between expected and observed logFC quickly increases with increasing developmental shifts up to 0.9, meaning that around 80% of the variance in logFCs is explained by development at 7 hours of time shift. At the same time, the performance of a standard linear model p-value in identifying truly DE genes quickly drops (Fig. 5f) as more and more expression changes are due to development (Sup. Fig. 24b).

We show that the strain logFC from the reference integrated model already performs better in detecting truly DE genes than the standard analysis p-value for developmental shifts of 3 or more hours (at 25°C, Sup. Fig. 24c). However, we propose an integrated predictor including the weighted mean between the standard analysis p-value and the strain logFC of the reference integrated where the optimal weight **w** is proportional to the variance of standard logFC explained by development (Sup. Fig. 24d, 24e). At the optimal **w** (see methods, Sup. Fig. 24e), our integrated predictor outperforms the standard analysis p-value for all time-shifts considered, with larger time-shifts showing the strongest improvements (Sup. Fig. 24c, 24d). For the largest shift (WT -7), **w** = 1 meaning that no information from the standard DE p-value is used to get the best results.

As no gold-standard is usually available to guide the choice of **w**, we explored the relationship between the optimal **w** and the correlation of logFCs with the reference. Sampling more WT subsets including non-contiguous sets (see methods, Sup. Table 8) reveals a tight relationship between optimal ***w*** and the correlation between observed and expected logFCs (Sup. Fig. 24f, 24g) which can therefore suggest the appropriate value of ***w***.

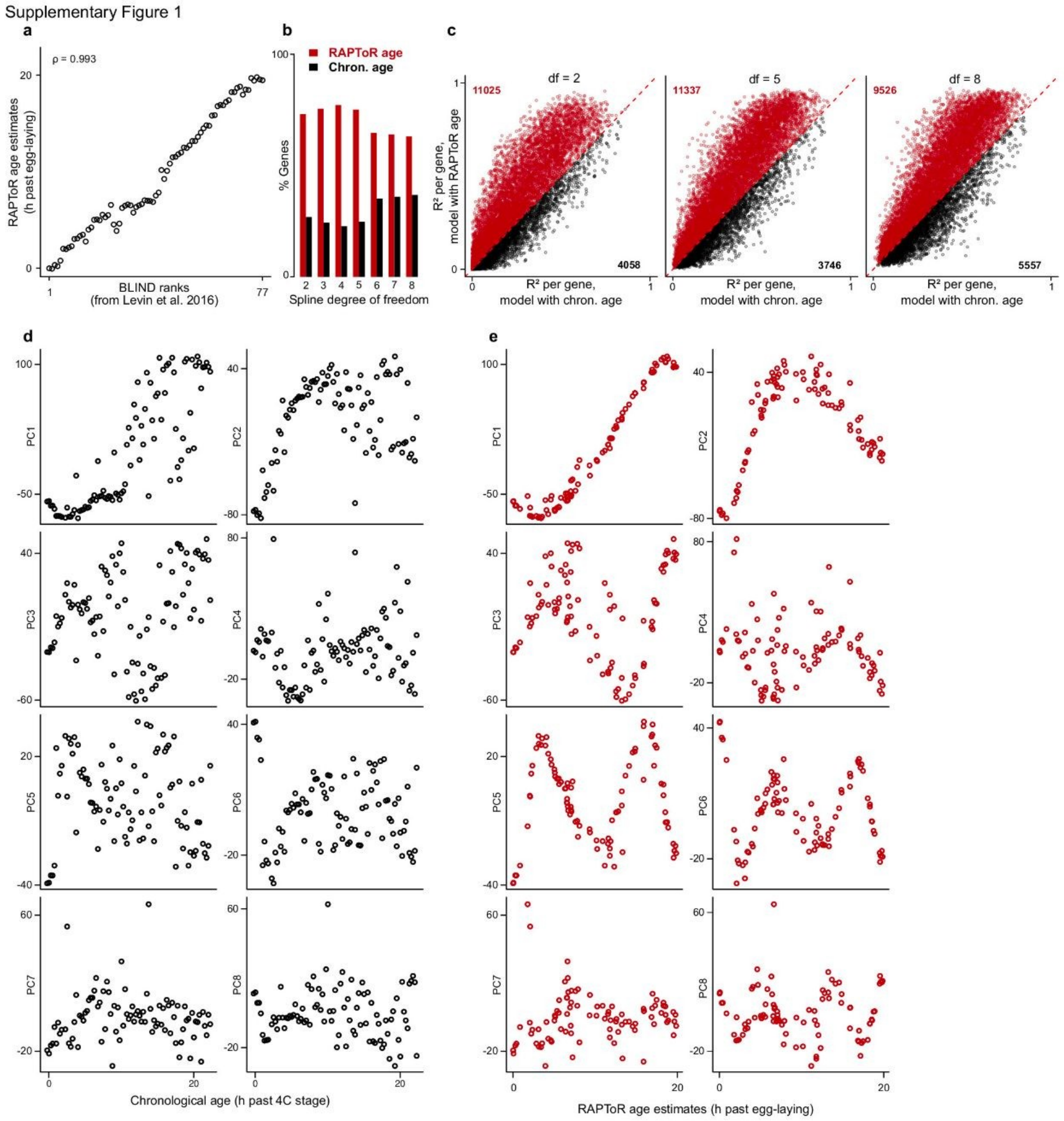


**Supplementary Figure 1 – RAPToR estimates fit gene expression data better than chronological age (related to Figure 2)**.

**a**, RAPToR estimates of *D. melanogaster* single-embryo samples[6] staged on a reference built from bulk data[16] plotted against established BLIND ranks[6] .

**b**, Percentage of genes better fitted by either RAPToR estimates or chronological age modeled with splines using from 2 to 8 degrees of freedom in otherwise identical models.

**c**, $R^2$ of models from (**b**) for. Gene count in each half of the plot is indicated in the corners.

**d,e**, Principal components plotted along chronological age (**d**), and RAPToR estimates (**e**) (as in Fig. 2d-f)

Supplementary Figure 2

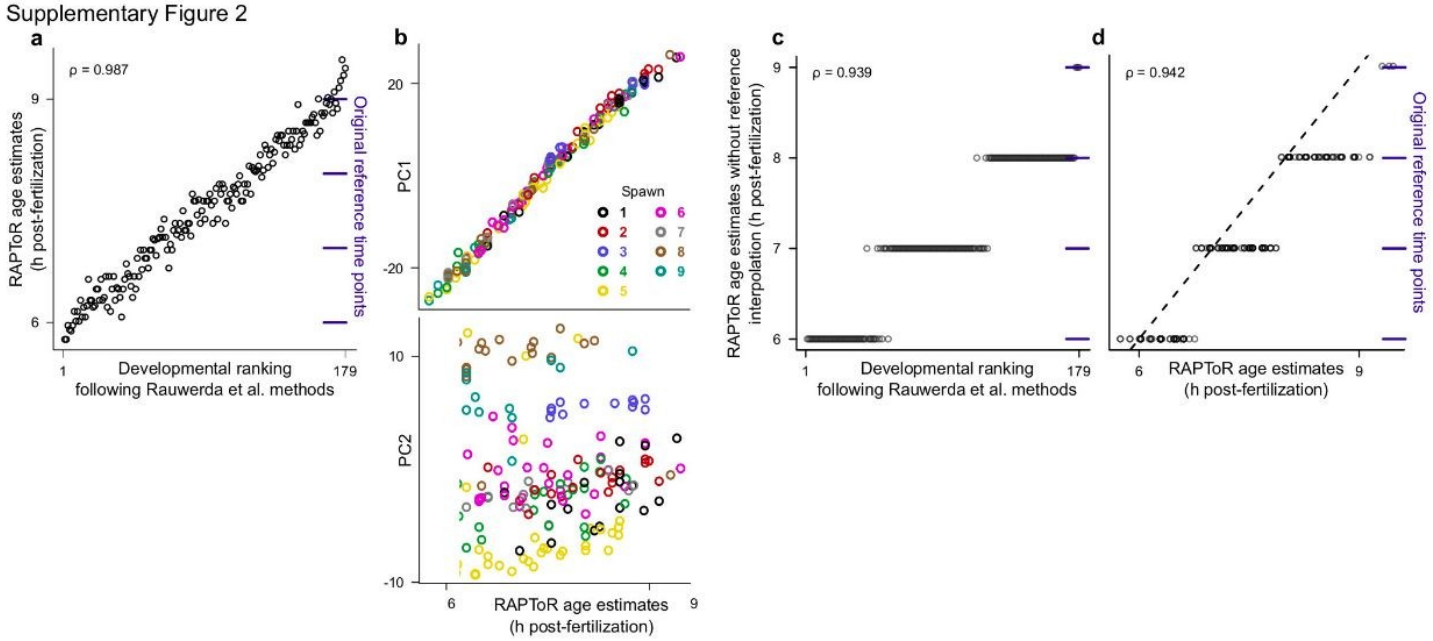


**Supplementary Figure 2 – Reference interpolation allows RAPToR estimates at high resolution**

**a**, RAPToR estimates of a zebrafish embryonic time series from 9 spawns[8] staged on a reference built from Domazet et al. data[10] plotted against original developmental ranks[8] .

**b**, First 2 principal components of the zebrafish time series from plotted against RAPToR age estimates. Spawns are color-coded.

**c,d**, RAPToR estimates of the zebrafish time series on the non-interpolated reference (i.e the sampling time of the reference sample with the highest correlation) vs. original developmental ranks (**c**) and vs. standard RAPToR estimates (as in **a)** (**d**).

In **a,c,d**, original reference time points within the plot area are shown on the right, in blue.

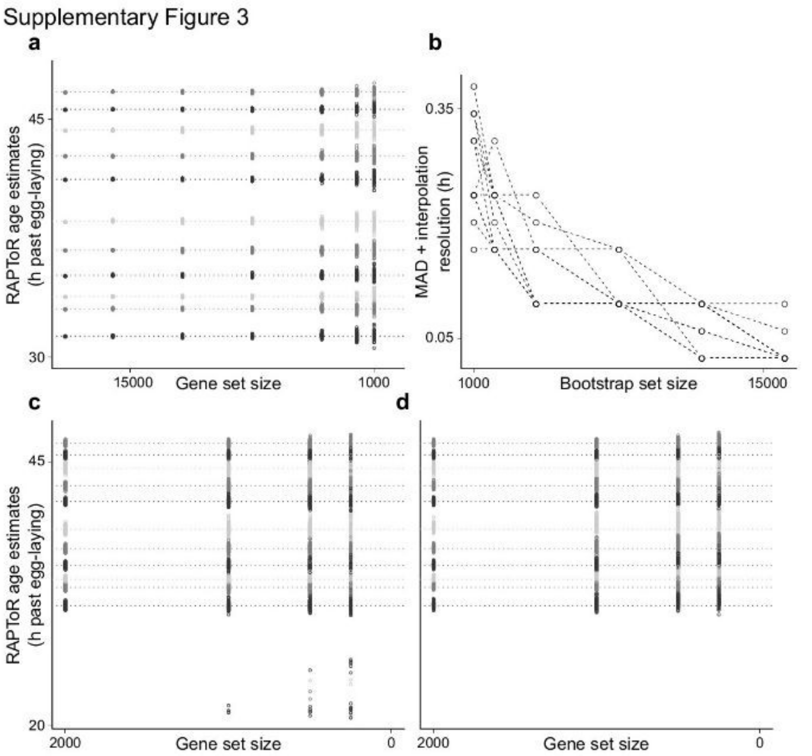


**Supplementary Figure 3 – Effect of gene set size on RAPToR estimates**

**a,** Effect of gene set size on RAPToR estimate of a *C. elegans* larval development time series[5] staged on a reference built from Kim et al.[9] data. Leftmost points and dashed lines indicate estimates on all genes (18718).

**b,** Median Absolute Deviation of bootstrap estimates from global estimate + interpolation resolution (i.e. half a confidence interval) by bootstrap set size for 50 bootstrap estimates.

**c,d,** Effect of small gene sets on RAPToR estimates without prior (**c**), or with prior (**d**). Some samples are staged to a previous molt (see also Sup. Fig. 4) when the gene set is small, an issue solved including a prior. Horizontal dashed lines indicate estimates of the samples using the full available gene set (18718).

Supplementary Figure 4

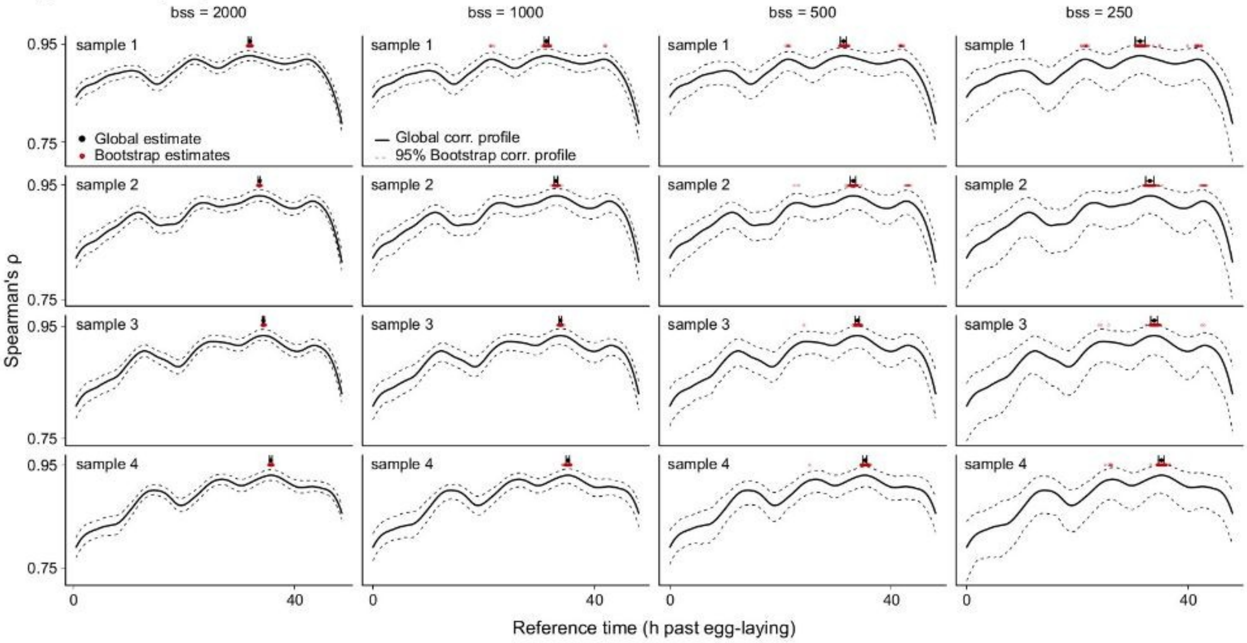


**Supplementary Figure 4 – Effect of bootstrap (or gene) set size on RAPToR correlation profiles**

Correlation profiles of the first 4 samples of the Hendriks et al.[5] time course staged on the reference built from Kim et al.[9] samples. Bootstrap gene set size (bss) was set to 2 000, 1 000, 500 or 250, with 100 bootstraps. With small gene sets, samples can be staged to other points in the transcriptomic landscape with similar expression (e.g, the four maxima in the profiles shown here are in phase with the oscillatory expression pattern of the four successive larval molts of *C. elegans*).

Global and bootstrap estimates, as well as the confidence interval, are shown above each profile. Dotted lines around the global correlation profiles correspond to 0.025 and 0.975 quantiles of the bootstrap correlation profiles. As expected, this interval gets larger for smaller bootstrap gene set sizes.

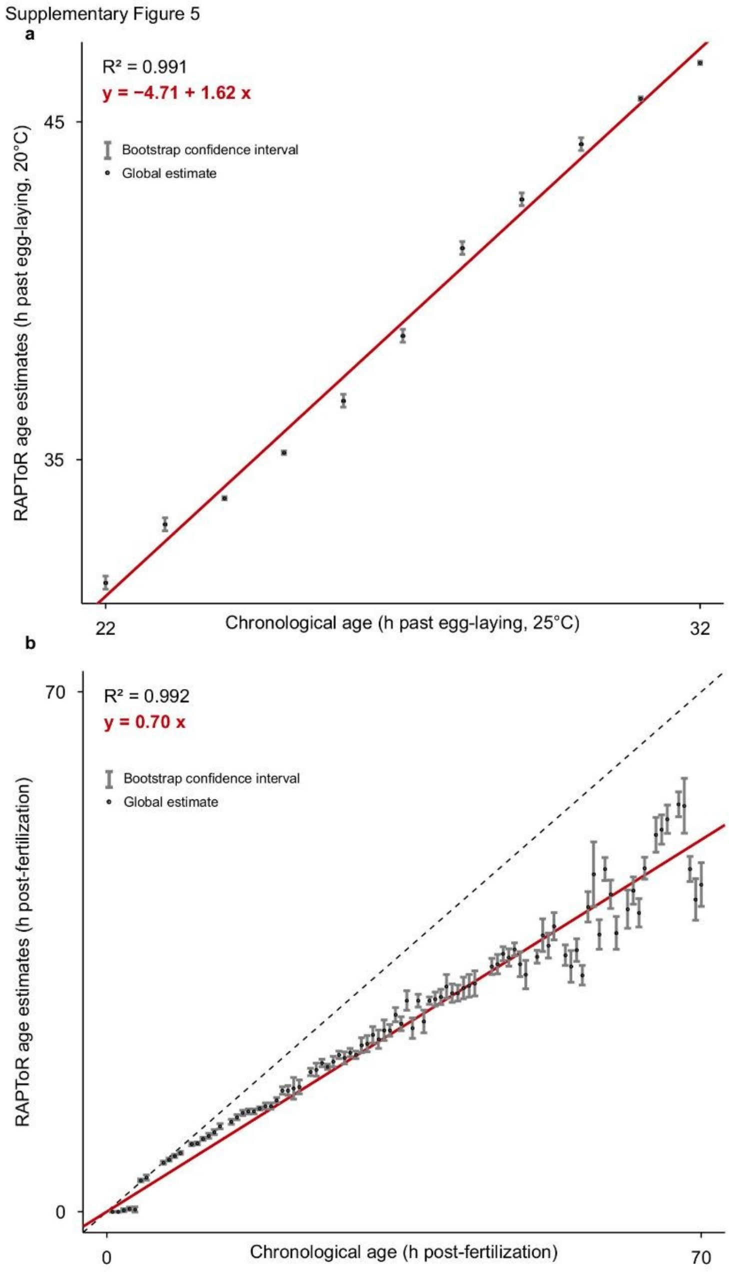


**Supplementary Figure 5 – Minute-scale precision staging of development**

**a,b**, Chronological age vs. RAPToR age estimates and their confidence intervals of *C. elegans* late larval development[5] (**a**), and *D. rerio* embryo development[6] (**b**) staged on appropriate references[9,10] (as in Fig 2a, 2b).

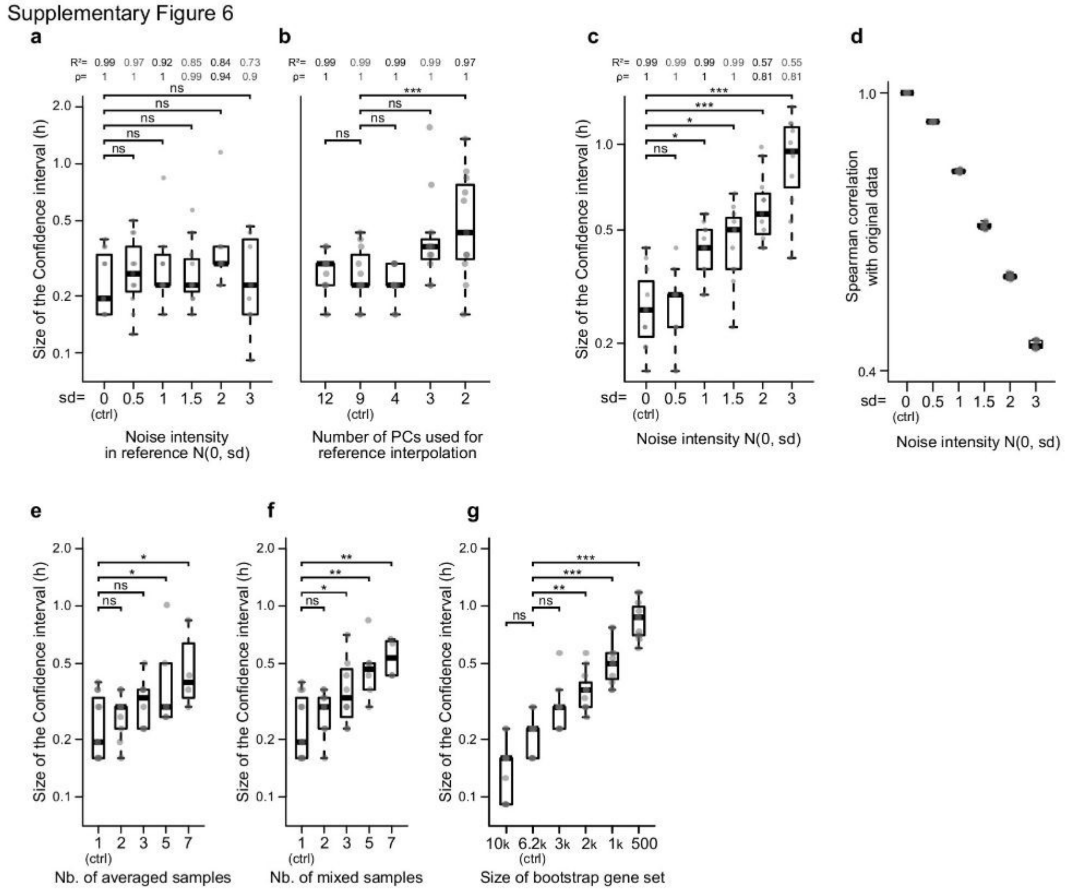


**Supplementary Figure 6 – Effect of data quality on staging and confidence intervals**

A *C. elegans* larval development time series[5] is staged on a reference built from an independent larval development time series[9]. Samples were staged using 10,000 genes, with bootstrap confidence intervals (CI) built from 50 bootstrap estimates.

**a,b**, Size of bootstrap CIs when adding increasing gaussian noise to log(TPM+1) reference data before interpolation (**a**), and when changing the number of components used for interpolation (**b**).
**c**, Size of bootstrap CIs when adding gaussian noise to log(TPM+1) sample expression data.
**d**, Spearman correlation between original data and samples + noise (as in **c**).
**e,f**, Size of bootstrap CIs when averaging samples together to mimic developmental spread of individuals (**e**) and when mixing samples together (randomly picking a gene expression value between n samples) to mimic heterochrony (**f**).

In **a-c**, $R^2$ and Spearman correlation between RAPToR age estimates and chronological age for each condition is displayed above the boxplots.
In **a-d**, each box is n=11; in **e,f**, boxes are n=11, 10, 9, 8, and 7 from left to right respectively.
In **a-c,e,f**, significance of mean difference with the control condition (noted “(ctrl)”) is tested with a linear model. P-values are adjusted for FDR within each panel. *: $p < 0.05$, **: $p < 0.01$, ***: $p < 0.001$.

Supplementary Figure 7

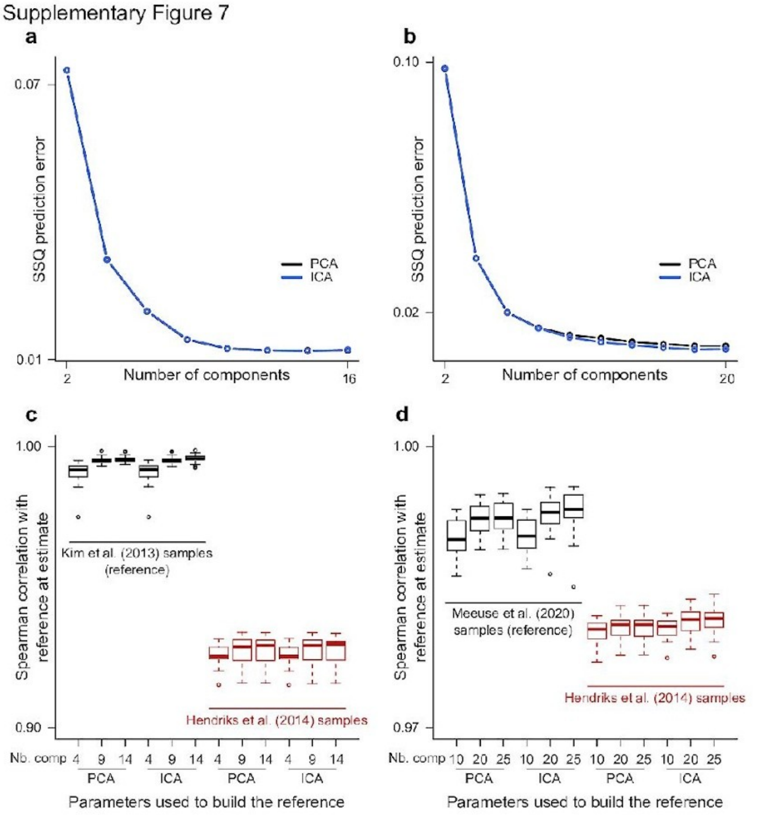


**Supplementary Figure 7 – Robustness of reference-building to parameter change**

**a,b**, Sum Squared prediction error of gene expression interpolation at known time points by number of components and dimension-reduction method used for interpolation in the Kim et al.[9] (**a**), and Meeuse et al.[15] (**b**) references of *C. elegans* larval development.

**c,d**, Spearman correlation between interpolated reference and (non-interpolated) reference and independent time series[5] samples at their age estimates when varying dimension reduction method and component number used for reference-building, using Kim et al. samples (**c**) or Meeuse et al. (**d**) samples to build a reference (see also Sup. Table 2).

In **c**, each box is n=26 for Kim et al.[9] samples and n=12 for Hendriks et al.[5] samples; in **d**, each box is n=44 for Meeuse et al.[15] samples and n=16 for Hendriks et al. samples.

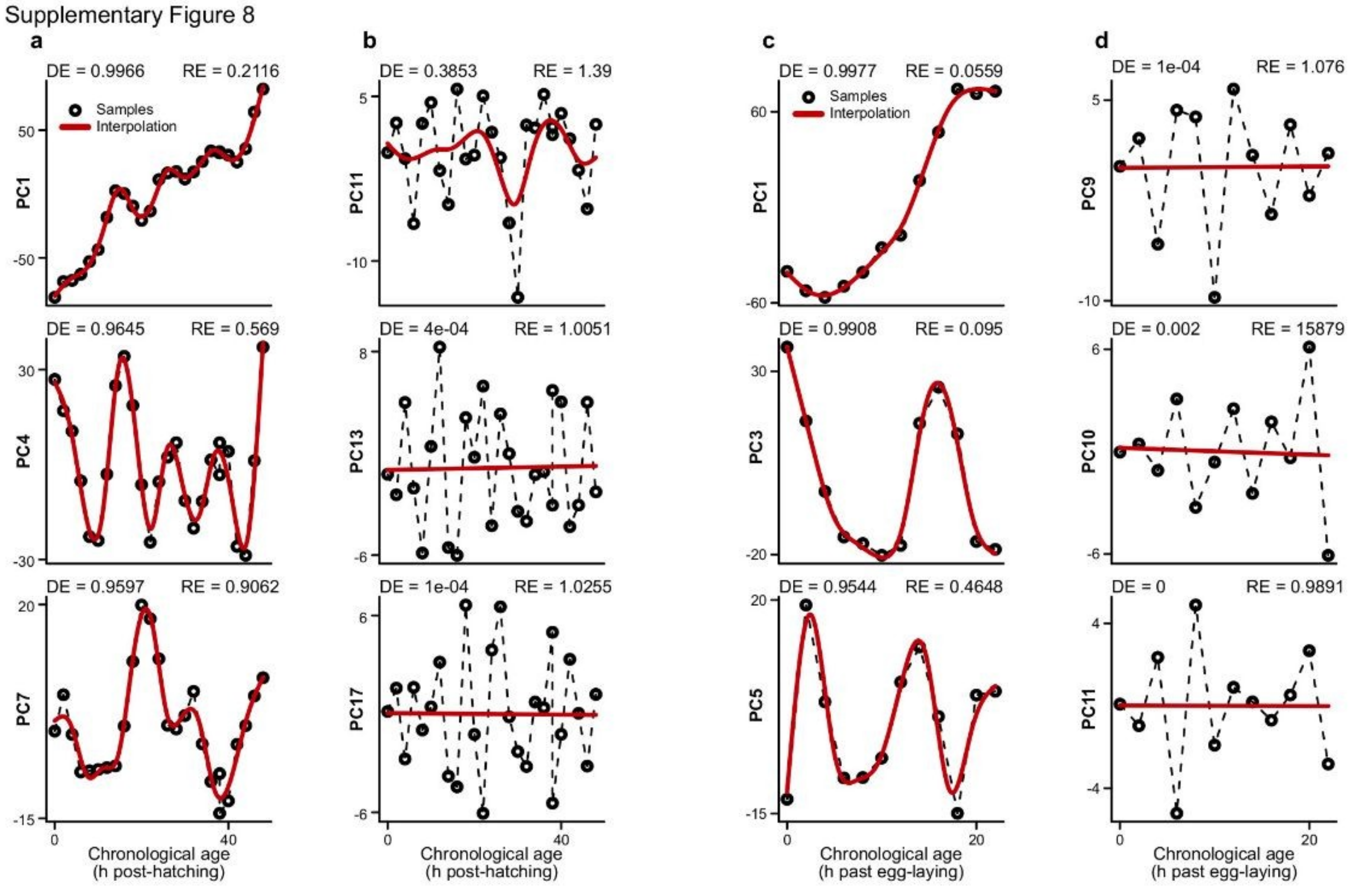


**Supplementary Figure 8 – Defining intelligible dynamics with spline fits**

Expression profiling time series are projected into principal component space and fit with their respective reference interpolation models (see Sup. Table 1). The interpolation of each component is then evaluated with the deviance explained (DE) and relative error (RE) of the fit.

**a,b**, *C. elegans* larval development[9] selected components with (**a**) and without (**b**) intelligible dynamics (with respect to time), that are kept or dropped for reference building respectively.
**c,d**, *D. melanogaster* embryo development[16] selected components with (**c**) and without (**d**) intelligible dynamics, that are kept or dropped for reference building respectively.

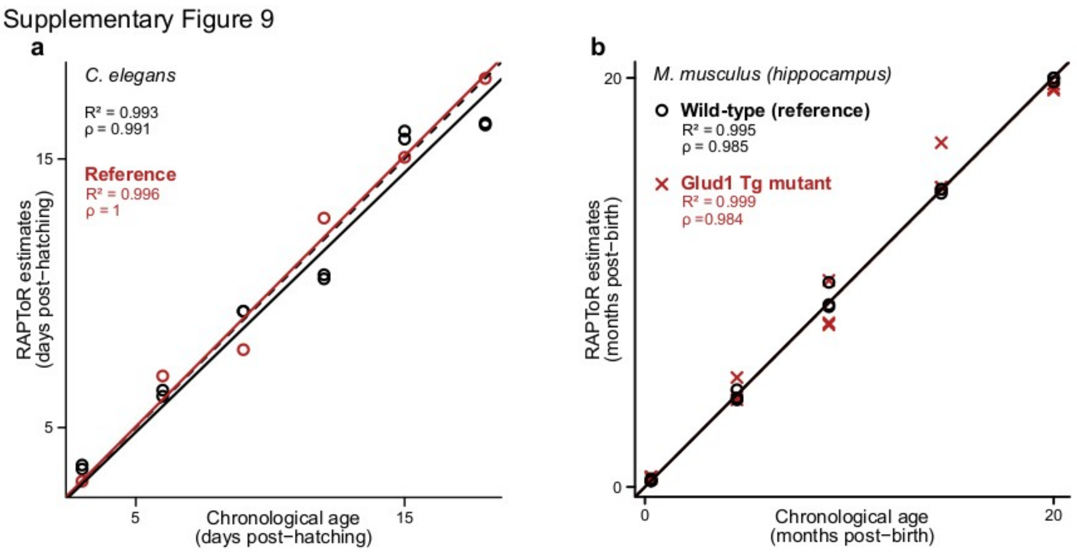


**Supplementary Figure 9 – RAPToR standard reference stage samples well within experiments**

**a**, Chronological age vs. RAPToR age estimates of a *C. elegans* aging time series (Byrne et al. 2020, unpublished). The samples from one of 3 replicates were used to build the reference (in red).

**b**, Chronological age vs. RAPToR age estimates of dissected hippocampus tissue from *M. musculus* across the entire lifespan of mice in wild-type (black) and *Glud1* transgenic (Tg) animals (red)[12]. Wild-type samples were used to build the reference.

Supplementary Figure 10

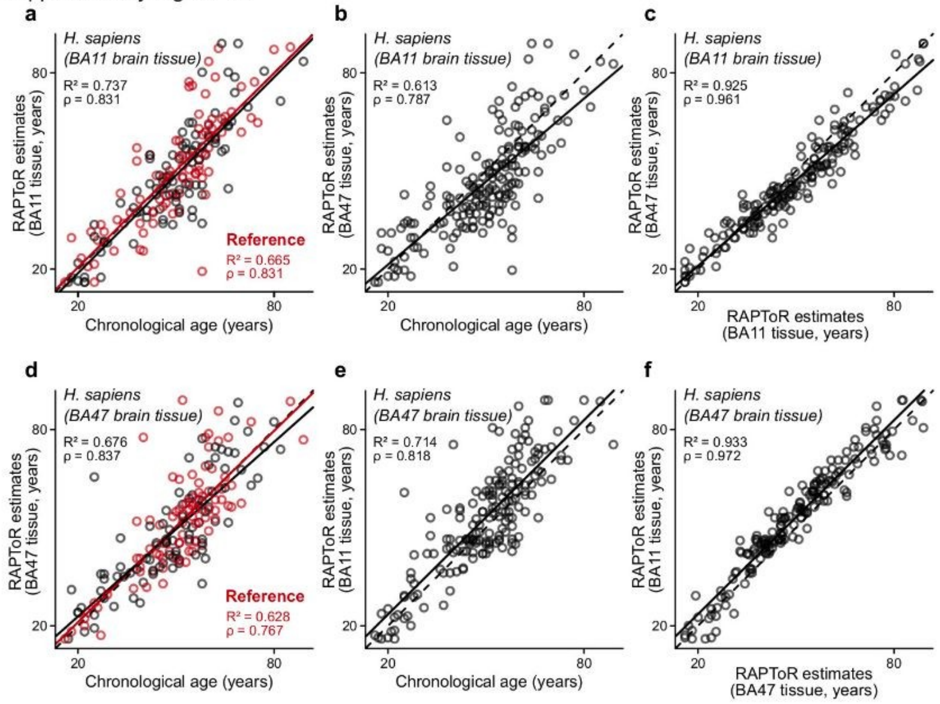


**Supplementary Figure 10 – RAPToR stages adult human brain tissue samples**

**a,b**, Chronological age vs. RAPToR estimates of human BA11 brain tissue samples on (**a**) a reference built from half the samples (in red) and (**b**) a reference built from human BA47 brain tissue samples.

**c**, RAPToR estimates of BA11 samples on BA11 reference (as in **a**) vs. RAPToR estimates on a BA47 reference (as in **b**).

**d,e**, Chronological age vs. RAPToR estimates of human BA47 brain tissue samples on (**d**) a reference built from half the samples (in red) and (**e**) a reference built from human BA11 brain tissue samples.

**f**, RAPToR estimates of BA47 samples on BA47 reference (as in **d**) vs. RAPToR estimates on a BA11 reference (as in **e**).

All samples are from Chen et al.[13]

Supplementary Figure 11

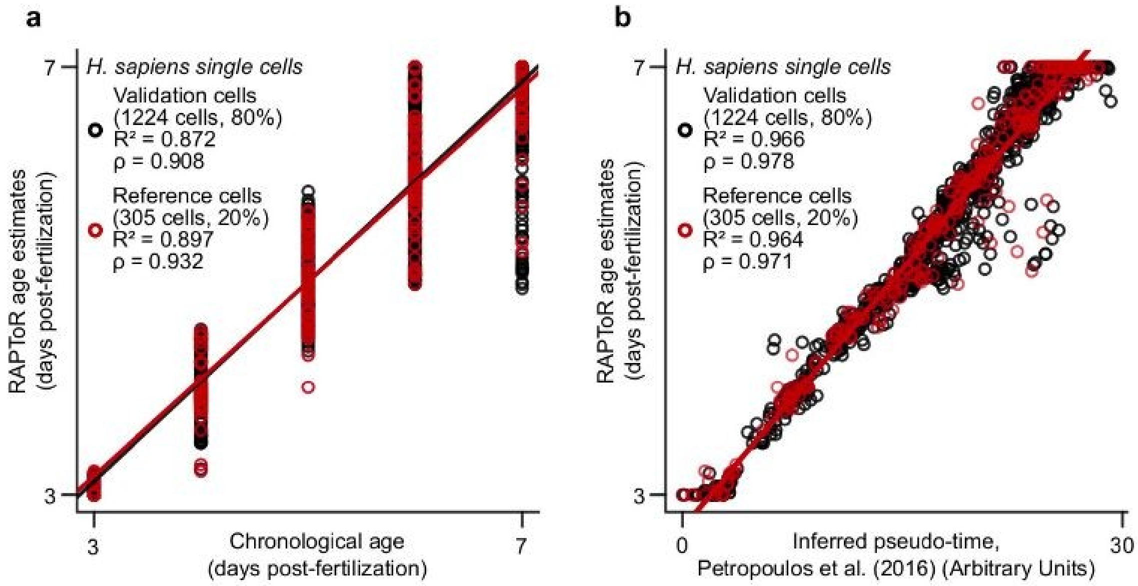


**Supplementary Figure 11 – Staging human single cells with RAPToR**

20% of a human early-embryogenesis single cell dataset[21] were used to build a reference, on which all cells were then staged. Metrics are split between reference and validation cell subsets.

**a**, Chronological age vs. RAPToR age estimates of single-cells.

**b**, Inferred pseudotime from the authors vs. RAPToR age estimates of single cells.

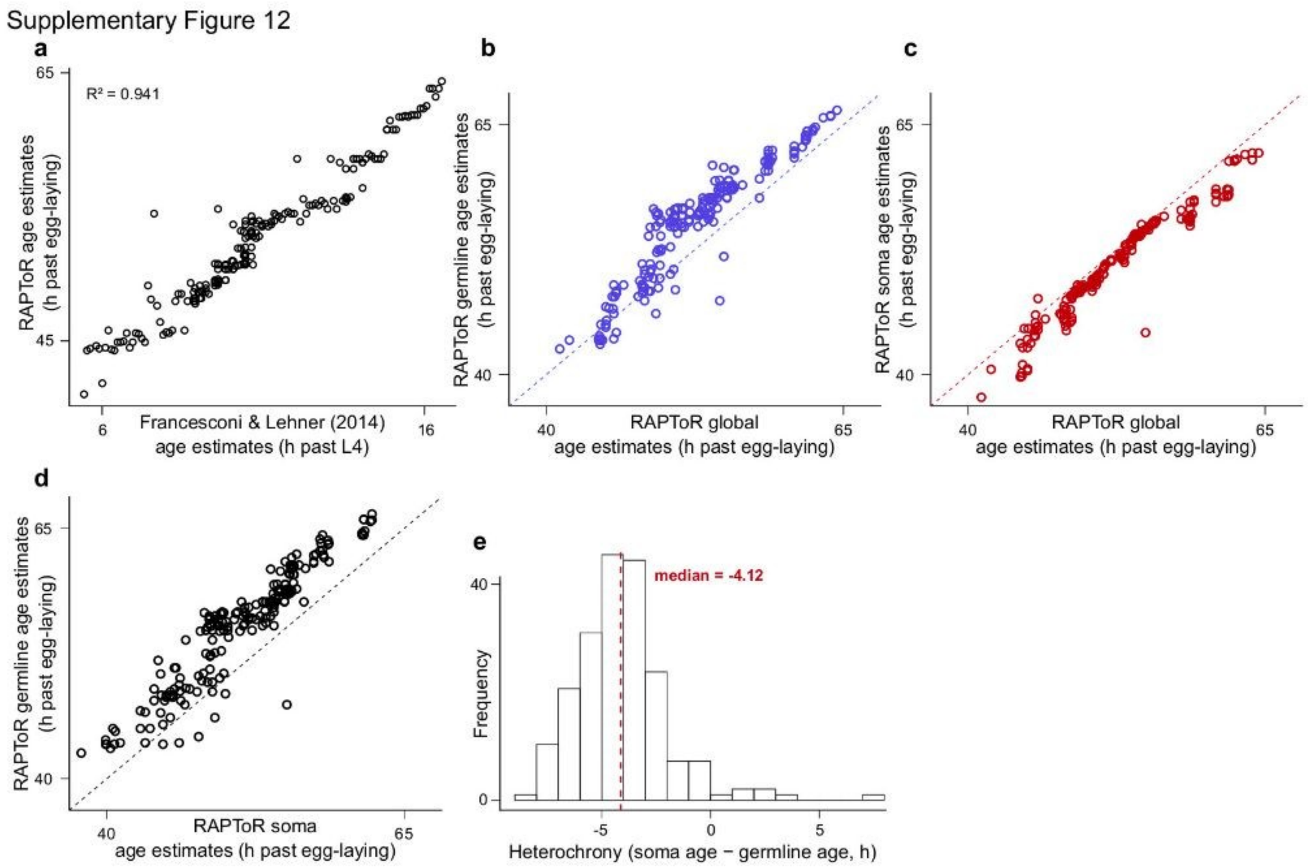


**Supplementary Figure 12 – Tissue-specific staging yields soma and germline ages**

**a**, RAPToR estimates of *C. elegans* Recombinant Inbred Lines (RILs)[14] staged on the larval to young-adult reference built from Meeuse et al.[15] vs. Francesconi & Lehner[22] estimates.

**b-d**, Comparison of RAPToR global age estimates vs. germline age estimates (**b**), global age estimates vs. soma age estimates (**c**), and soma age estimates vs. germline age estimates (**d**).

**e**, Distribution of soma-germline heterochrony.

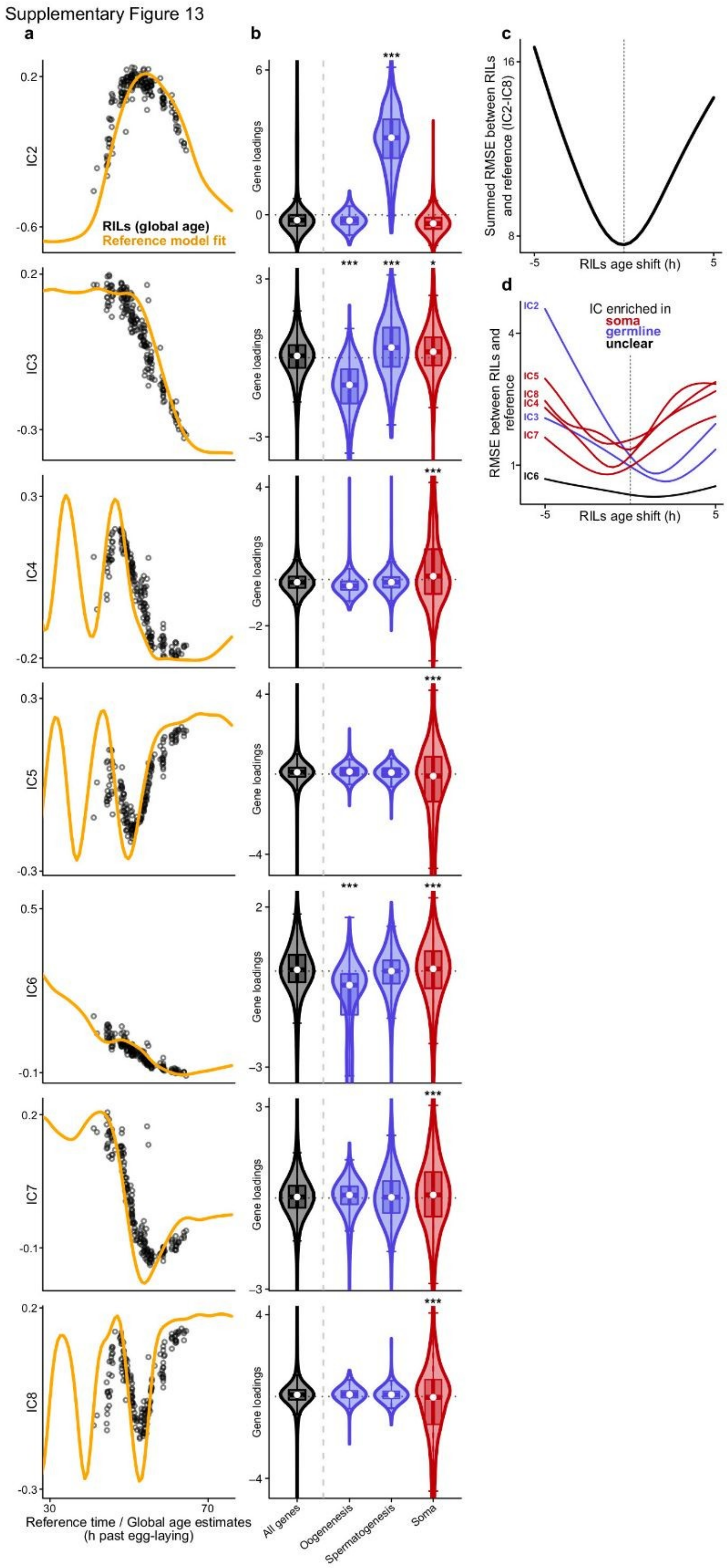


**Supplementary Figure 13 – A delayed germline and an advanced soma**

**a**, ICA components from ICA on *C. elegans* Recombinant Inbred Lines (RILs)[14] joined to the (non interpolated) reference data[15] plotted along chronological age and RAPToR global estimates for the reference (orange) and RILs (black) respectively.

**b**, Enrichment of gene loadings on ICA components for germline and soma categories.

**c,d**, Summed (**c**) and per-component (**d**) RMSE between RILs and reference fit on IC2-IC8 when shifting RIL (global) age estimates. RMSE per-component shows heterochrony, with soma dynamics of RILs matching younger reference time and the reverse for germline dynamics.

*: p < 0.05, **: p < 0.01, ***: p < 0.001.

RMSE: Root Mean Square Error

Supplementary Figure 14

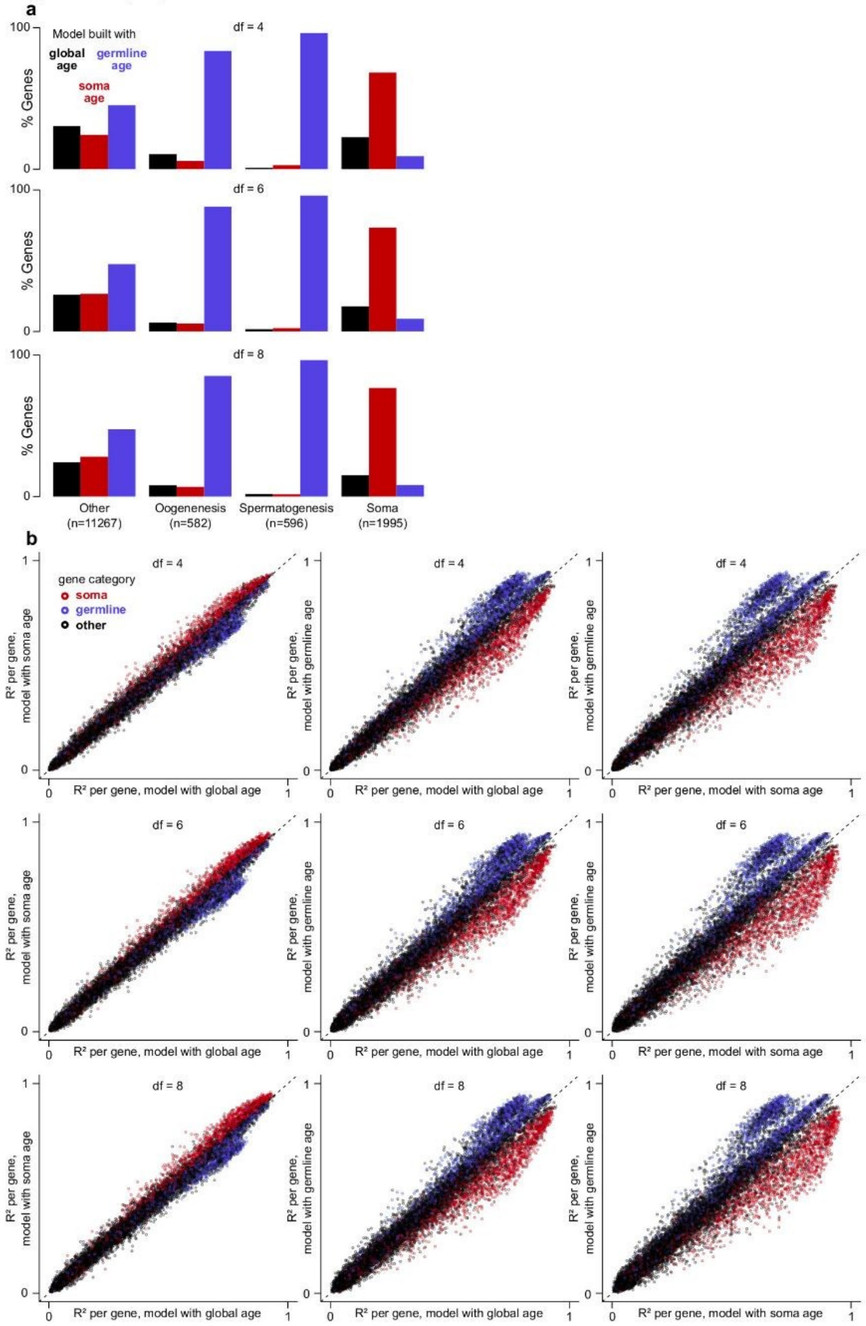


**Supplementary Figure 14 – Soma-germline heterochrony among *C. elegans* recombinant lines.**

Recombinant Inbred Lines (RILs)[14] are staged on the larval to young-adult reference built from Meeuse et al. samples[15] .

**a**, Percentage of genes better fitted by either RAPToR global, soma, or germline age estimates, modeled with splines with 4, 6 or 8 degrees of freedom in otherwise identical models. Genes are classified into spermatogenesis, oogenesis, somatic or other (see methods).

**b**, $R^2$ per gene of models with global, soma or germline age estimates as predictors for 4, 6,and 8 spline degree of freedom values.

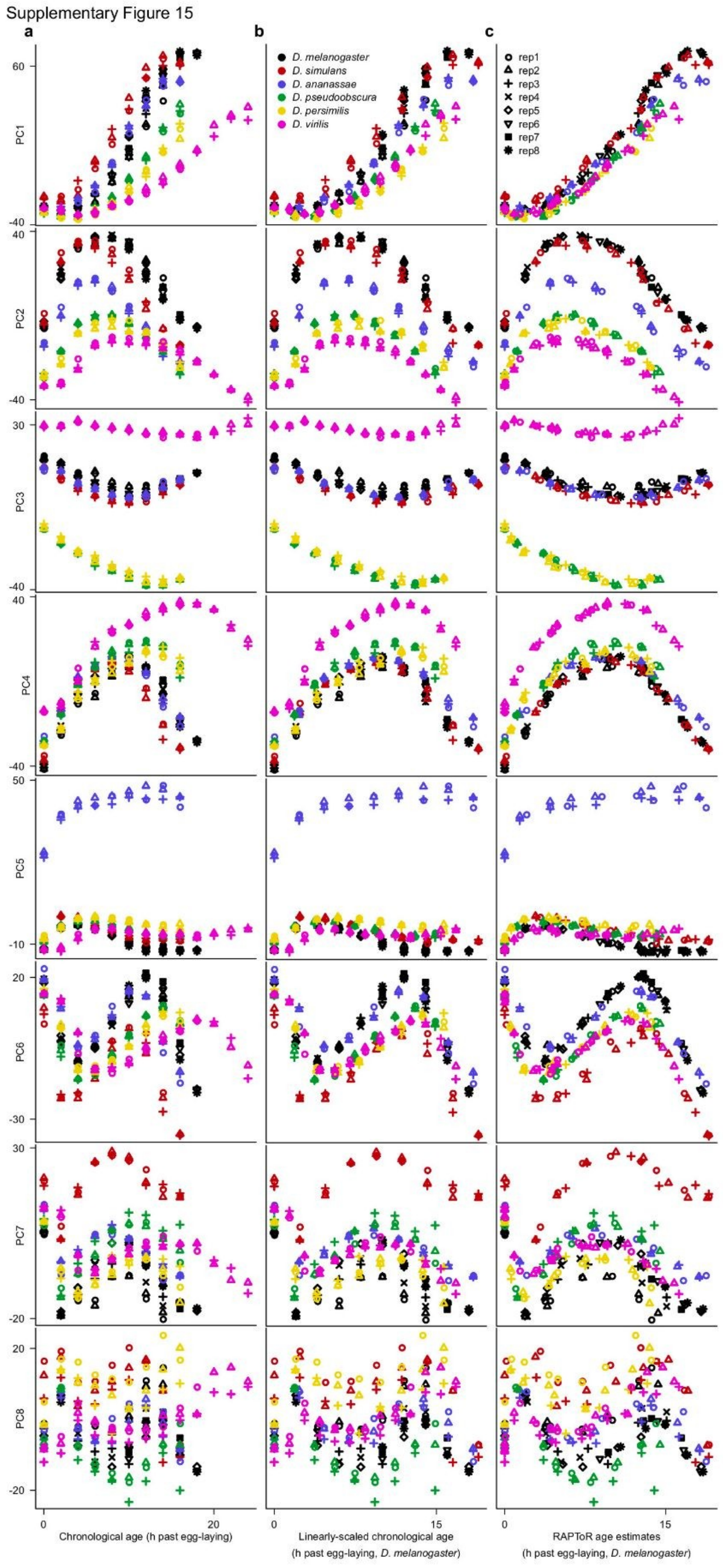


**Supplementary Figure 15 – RAPToR age estimates synchronize expression dynamics across species**

**a-c**, Principal components of *Drosophila* embryogenesis in 6 species[23] plotted along chronological age (**a**), linearly-scaled chronological age[23] (**b**), and RAPToR age estimates on a *D. melanogaster* reference[16] (**c**).

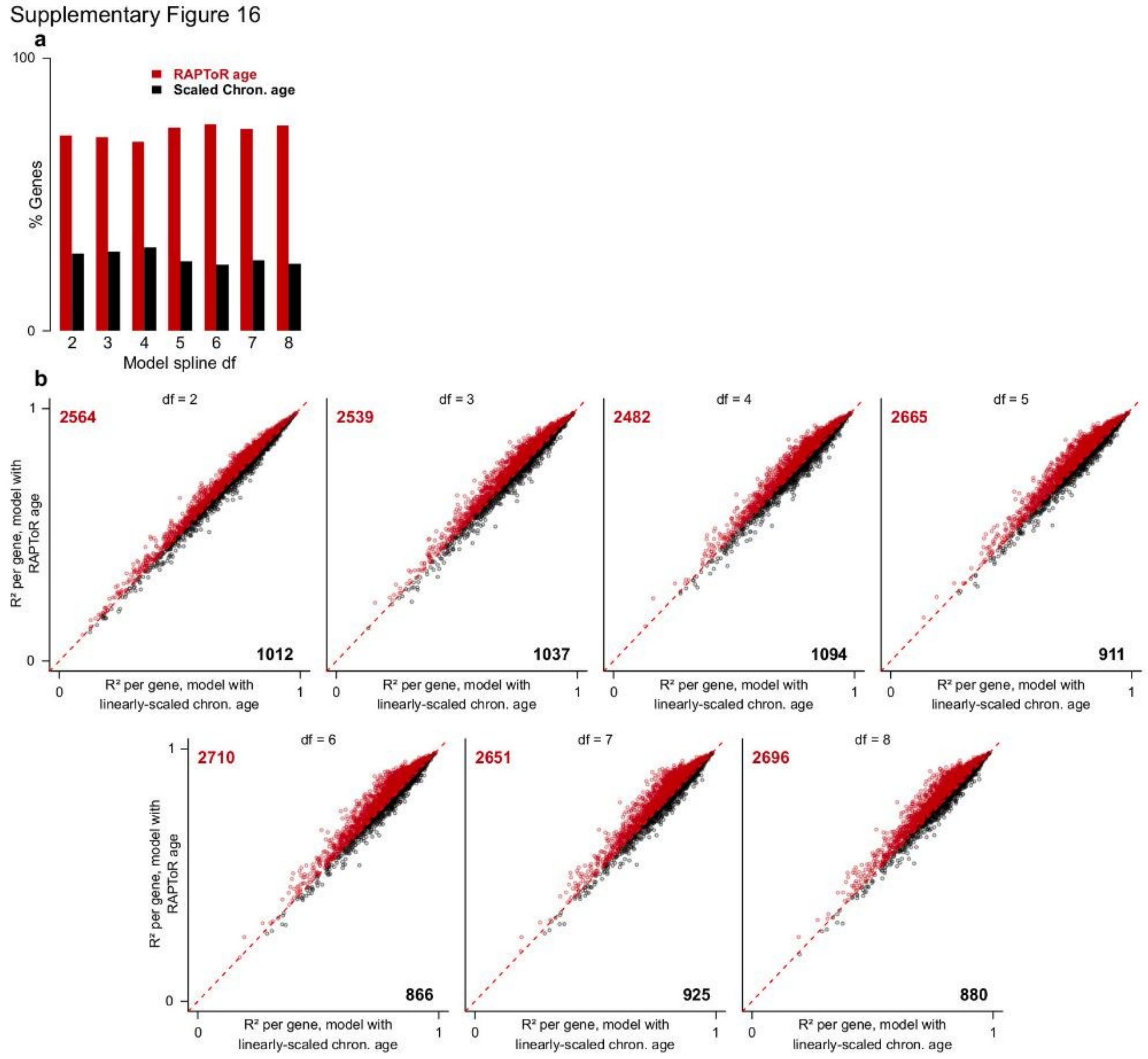


**Supplementary Figure 16 – RAPToR age estimates improve model fits over linear age scaling**
*Drosophila e*mbryo samples from Kalinka et al.[23] of 6 species are staged on a *D. melanogaster* reference built from Graveley et al.[16] samples (as in Fig 3a).

**a**, Choice between identical models fit on gene expression with linearly-scaled chronological age or RAPToR estimates as predictors.

**b**, $R^2$ per gene of models with chronological age as predictor vs. the same model using RAPToR estimates, across 2-8 of spline degrees of freedom.

Supplementary Figure 17

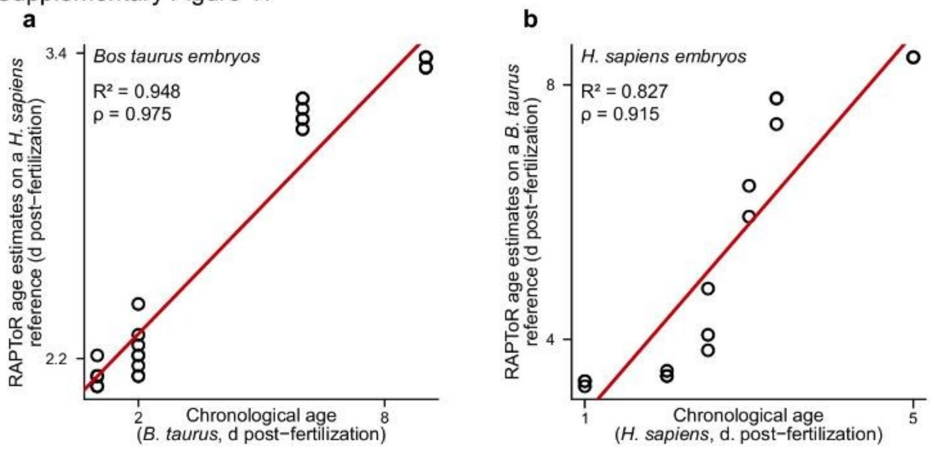


**Supplementary Figure 17 – Staging cow embryos on human embryos and vice versa**

**a**, Staging early-embryo development of *B. taurus*[24] on a human early-embryo development reference[25].

**b**, Staging early-embryo development of *H. sapiens*[25] on a cow early-embryo development reference[24].

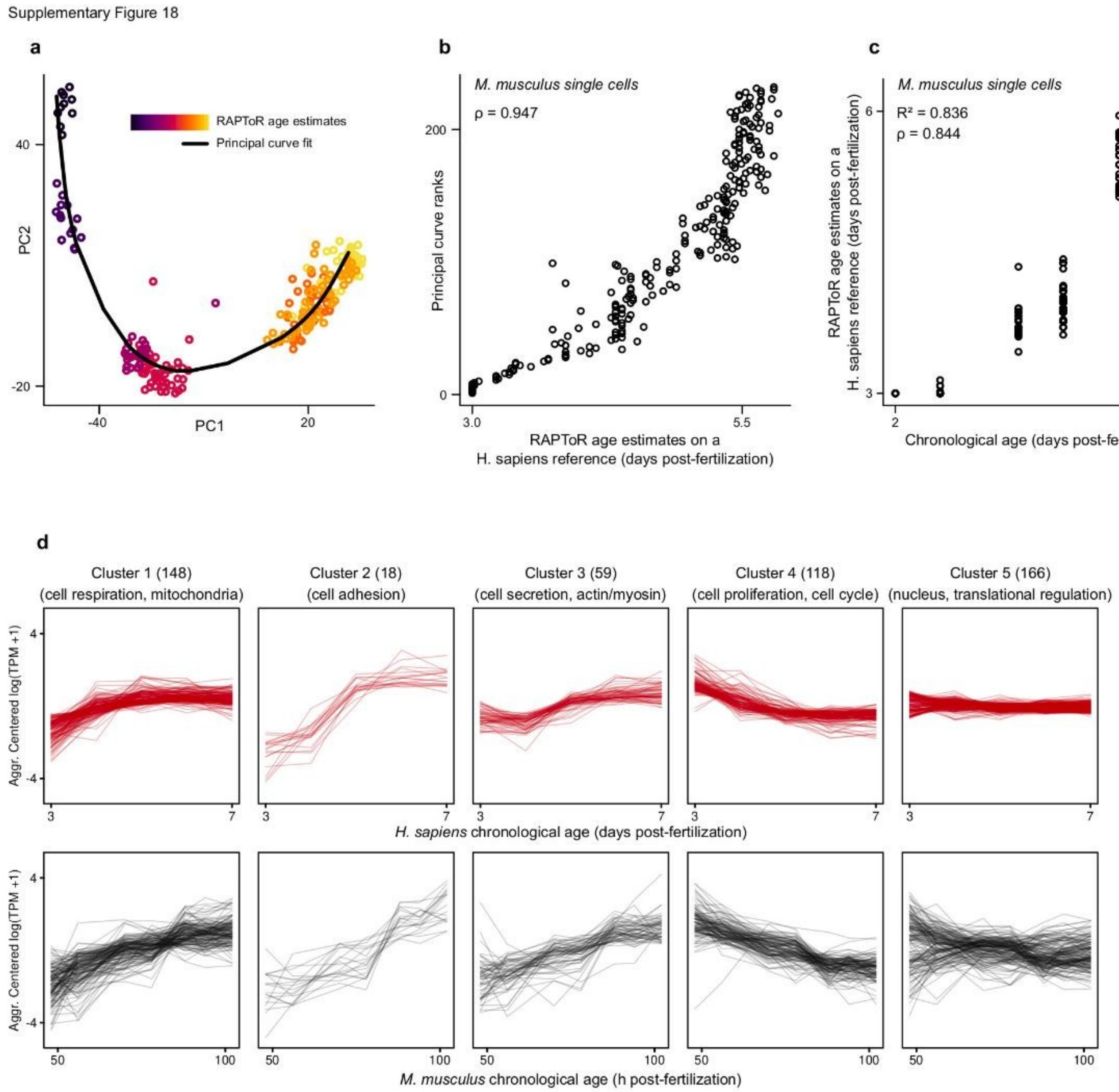


**Supplementary Figure 18 – Staging M. musculus single cells on H. sapiens reference**

Single cells from *M. musculus* embryos[26] were staged on a *H. sapiens* single-cell embryogenesis reference[21] using orthologs.

**a**, First 2 principal components of a PCA done on the 1000 most variable genes. A principal curve is fit on the first 3 components. Cells are colored by RAPToR age estimate on the *H. sapiens* reference.
**b**, RAPToR age estimates of *M. musculus* single cells on *H. sapiens* reference vs. cell ranks along principal curve (a).
**c**, Chronological age of *M. musculus* single cells vs. RAPToR age estimates on *H. sapiens* reference using top 10% most correlated genes for staging (see methods).
**d**, *H. sapiens* (red) and *M. musculus* (black) clustered gene expression profiles (aggregated per time point) of highest-correlated genes between both species (see methods).

Supplementary Figure 19

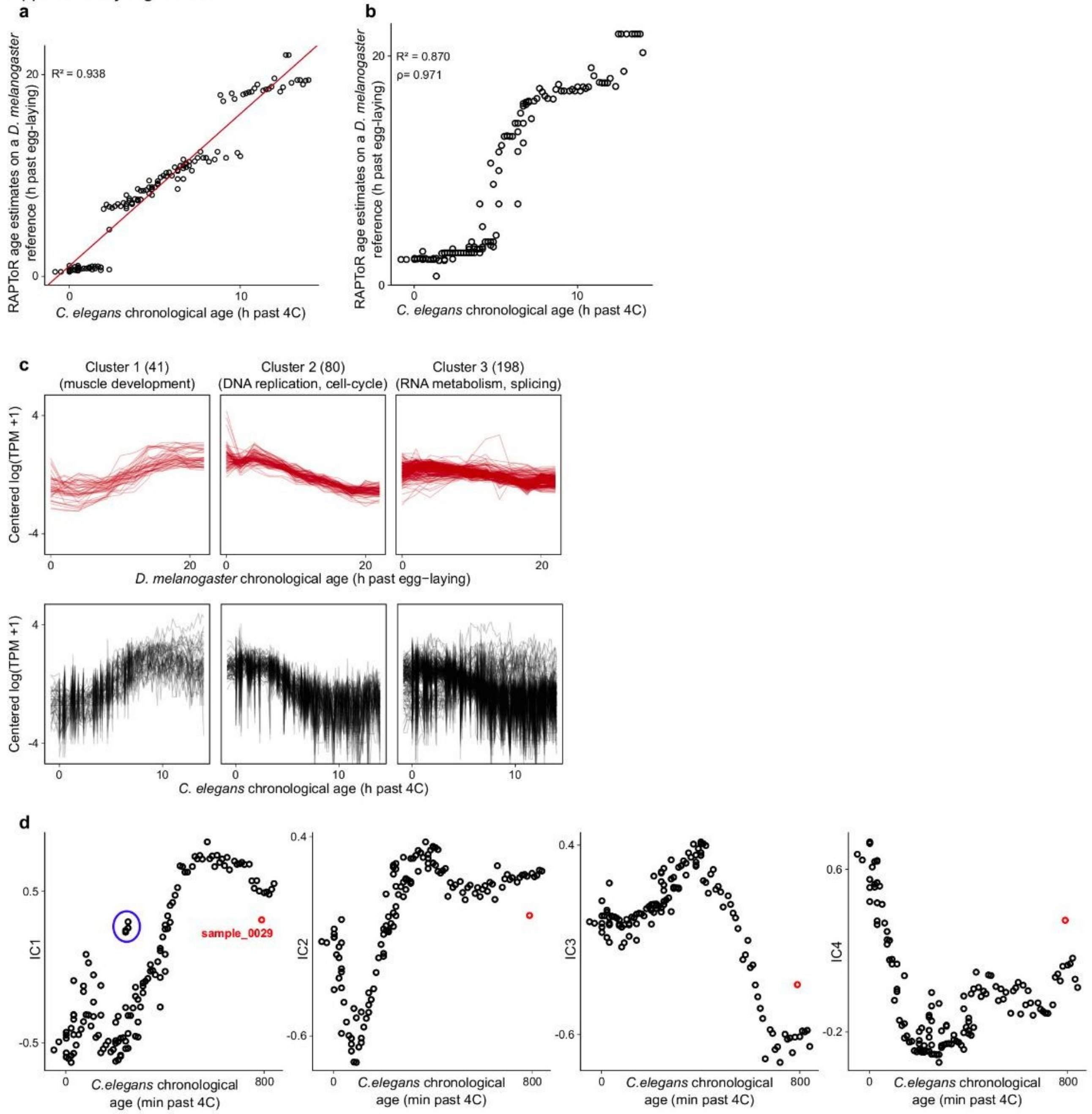


**Supplementary Figure 19 – Staging *C. elegans* embryogenesis with *D. melanogaster***

**a,** *C. elegans* embryo samples from Levin et al.[6] staged on the *D. melanogaster* reference built from Graveley et al.[16] samples. Gaps appear in the estimates, likely at points where fly expression dynamics are incompatible with those of worms.

**b,** As in (**a**), staging on the adjusted fly reference and using only highest-correlated genes between fly and worm embryogenesis (see methods).

**c,** *D. melanogaster* (red) and *C. elegans* (black) clustered gene expression profiles of highest-correlated genes between both species (see methods).

**d**, ICA components of the *C. elegans* embryo time course plotted along sampling time. Both the red highlighted outlier and 4 samples with erroneous chronological age (circled in IC1) are omitted from analysis (see methods).

Supplementary Figure 20

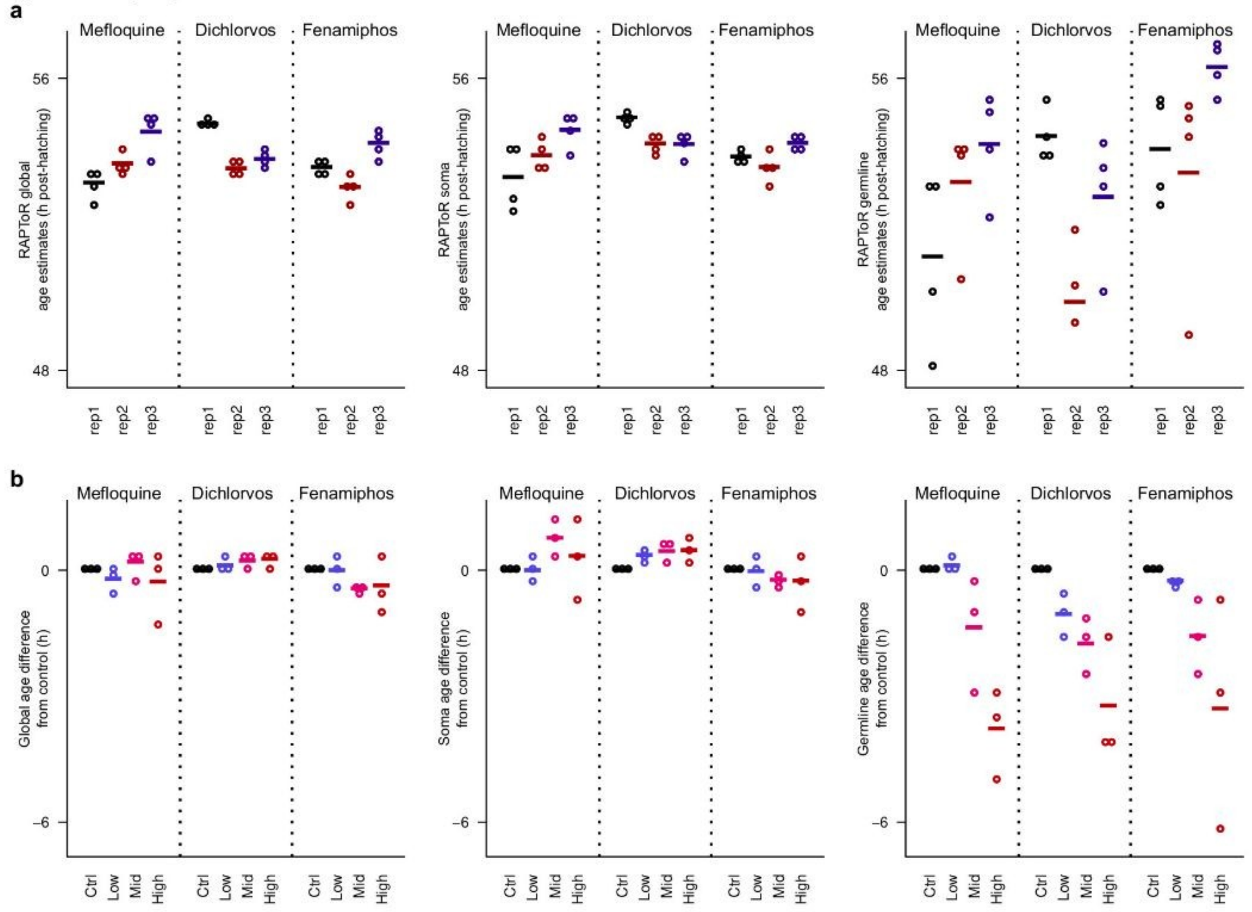


**Supplementary Figure 20 – Impact of drugs on C. elegans germline development**

*C. elegans* samples exposed to 3 doses of 3 different drugs profiled by Lewis et al.[17] are staged on the reference built from Meeuse et al.[15] samples.

**a,** Impact of batch on global, soma and germline age.

**b,** Impact of drug dose on global, soma and germline age, normalized per batch. Age difference is computed by subtracting the age of the control sample within each batch.

In **a,b**, bars indicate group mean.

Supplementary Figure 21

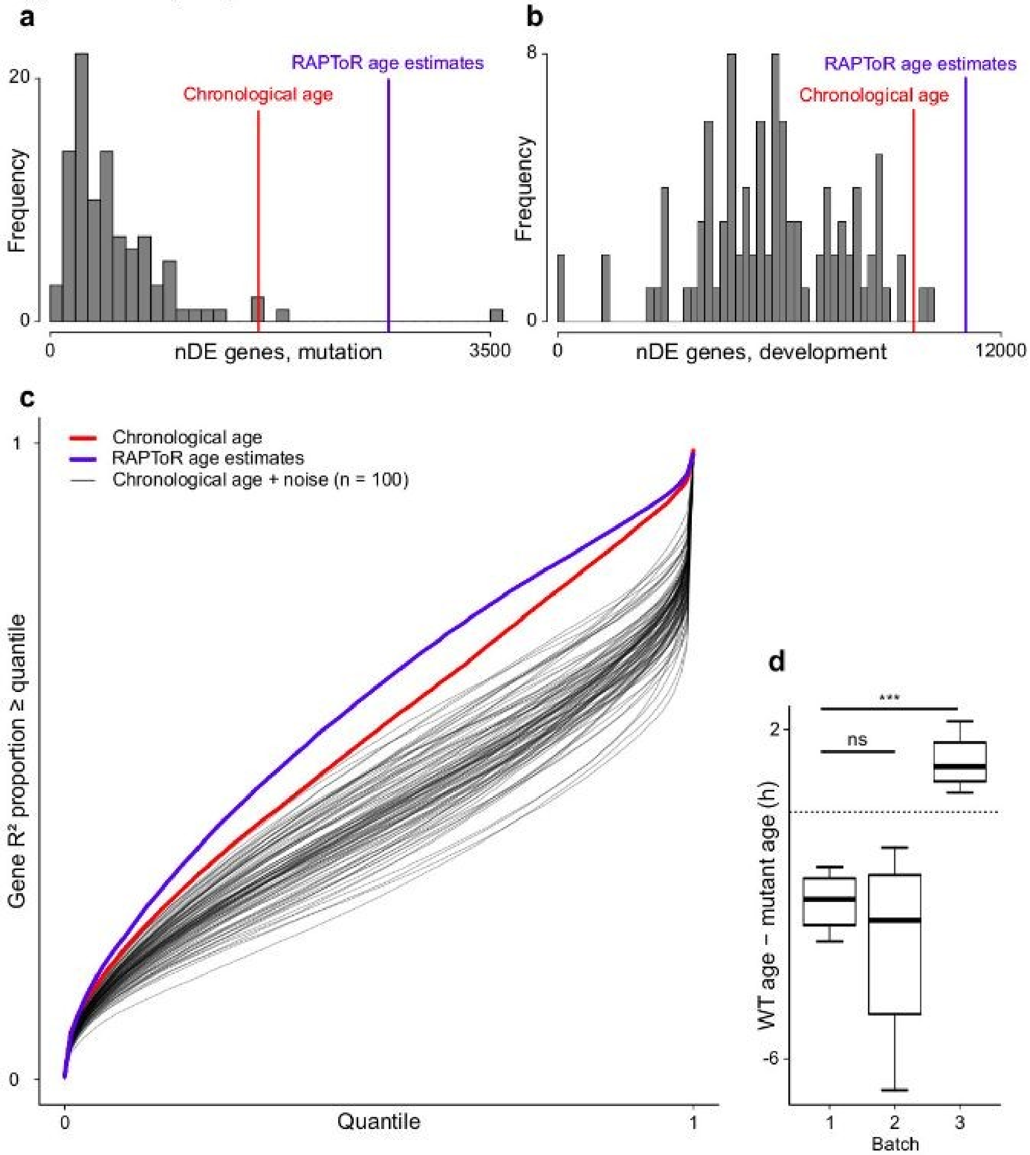


**Supplementary Figure 21 – Increasing the power of DE gene detection with RAPToR estimates**

*pash-1*ts or wild-type C. *elegans* samples profiled by Lehrbach et al.[18] are staged on the reference built from Reinke et al.[4] samples.

**a-c,** Random perturbations on chronological age (n=100 sets, see methods) produce overall fewer DE genes for strain (**a**), and development (**b**) than either chronological age (red) or RAPToR estimates (blue). These random perturbations further cause poorer model fits than chronological age whereas RAPToR age estimates systematically outperform chronological age (**c**).

**d**, Batch effect on developmental difference between control and *pash-1*ts samples. Each box is n = 4.

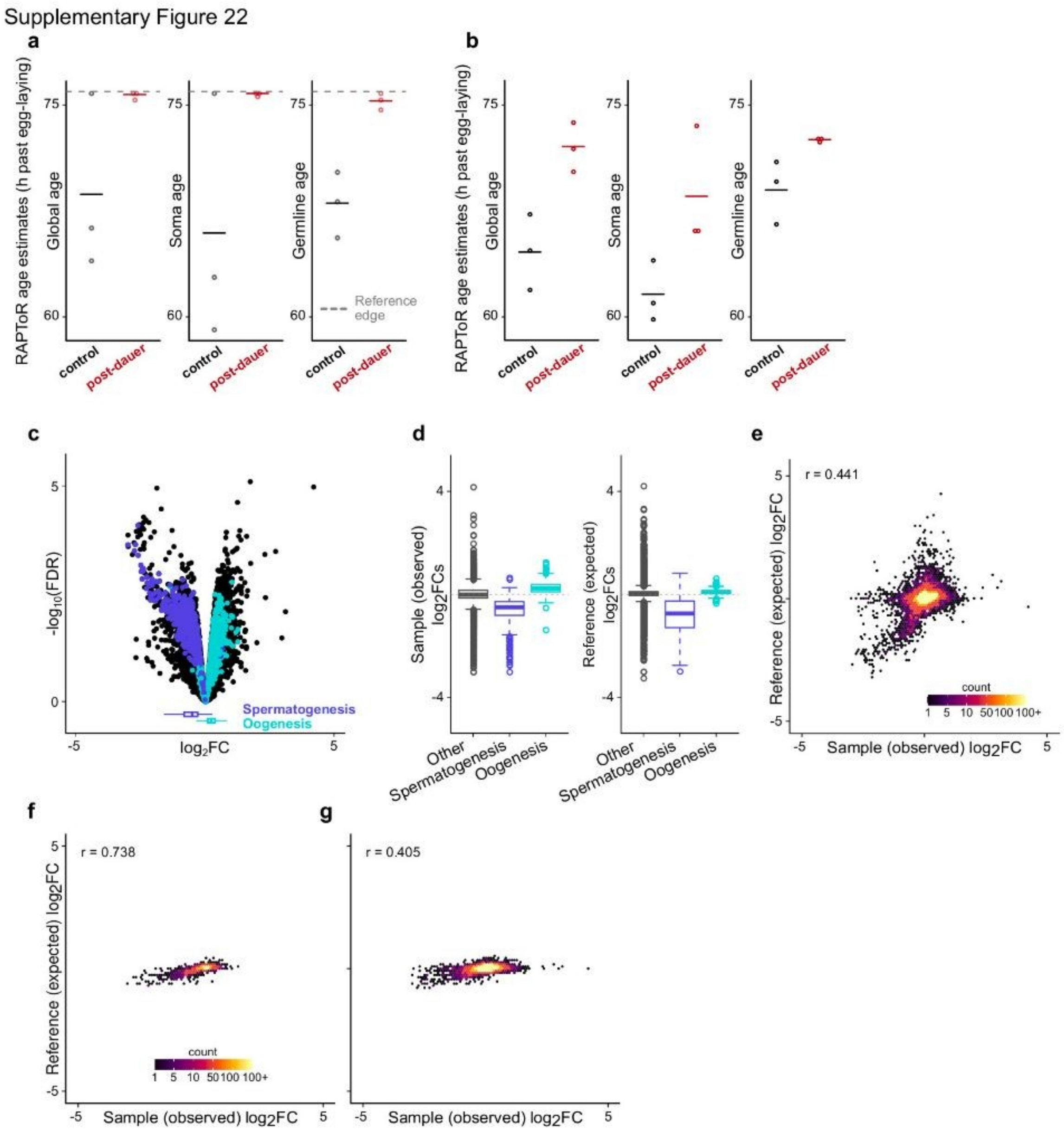


**Supplementary Figure 22 – Germline expression changes recapitulated by a developmental shift**

**a,b**, Global, soma-specific and germline-specific RAPToR age estimates of *C. elegans* control and post-dauer (PD) samples[19] staged on a reference built from Meeuse et al.[15] (**a**), and Reinke et al.[4] (**b**). Bars indicate mean age per group.

**c**, Volcanoplot of control vs. PD groups. Spermatogenesis and oogenesis genes are color-coded and logFCs of both categories are shown in boxplots at the bottom.

**d**, logFCs of control vs. PD for spermatogenesis and oogenesis genes observed in samples (left) and expected from development in the reference built from Meeuse et al., (right)

**d,** Observed expression logFCs between control and PD samples vs. expected developmental expression logFCs from the reference built from Meeuse et al. (as in Fig 5h, but for all genes)

**e,f,** Observed expression logFCs genes between control and PD samples vs. expected developmental expression logFCs from the reference built from Reinke et al., for germline genes (**e**), and all genes (**f**).

logFC: log2 fold-change, FDR: false discovery rate.

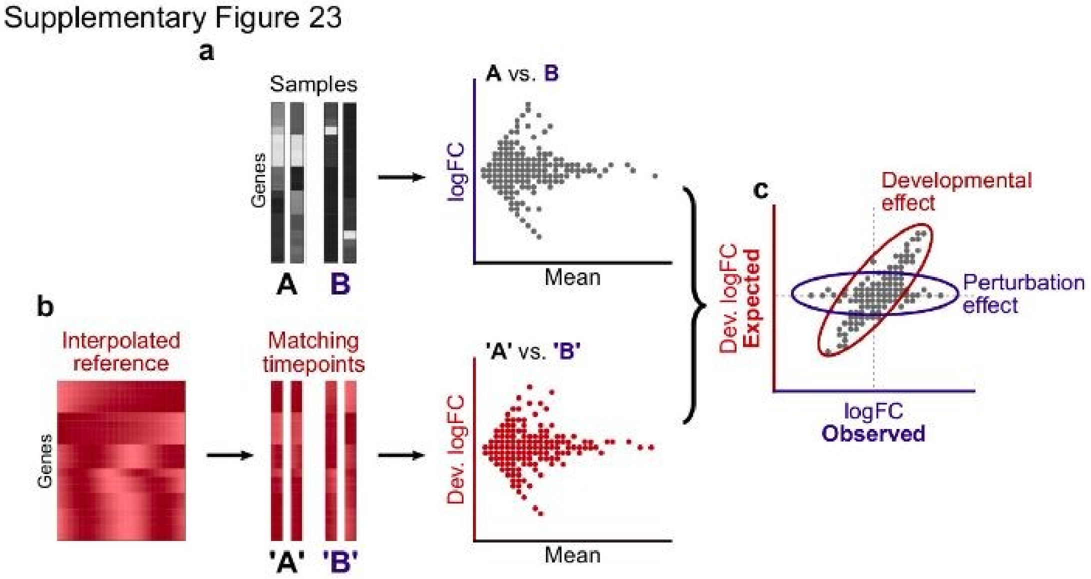


**Supplementary Figure 23 – Estimating the impact of development by integrating reference data**

**a-c**, Cartoon detailing how the logFCs of a differential expression analysis between two sample groups (**a**) and the logFCs of their matching time points in the RAPToR interpolated reference (**b**) can be compared to quantify the impact of development (**c**).

Supplementary Figure 24

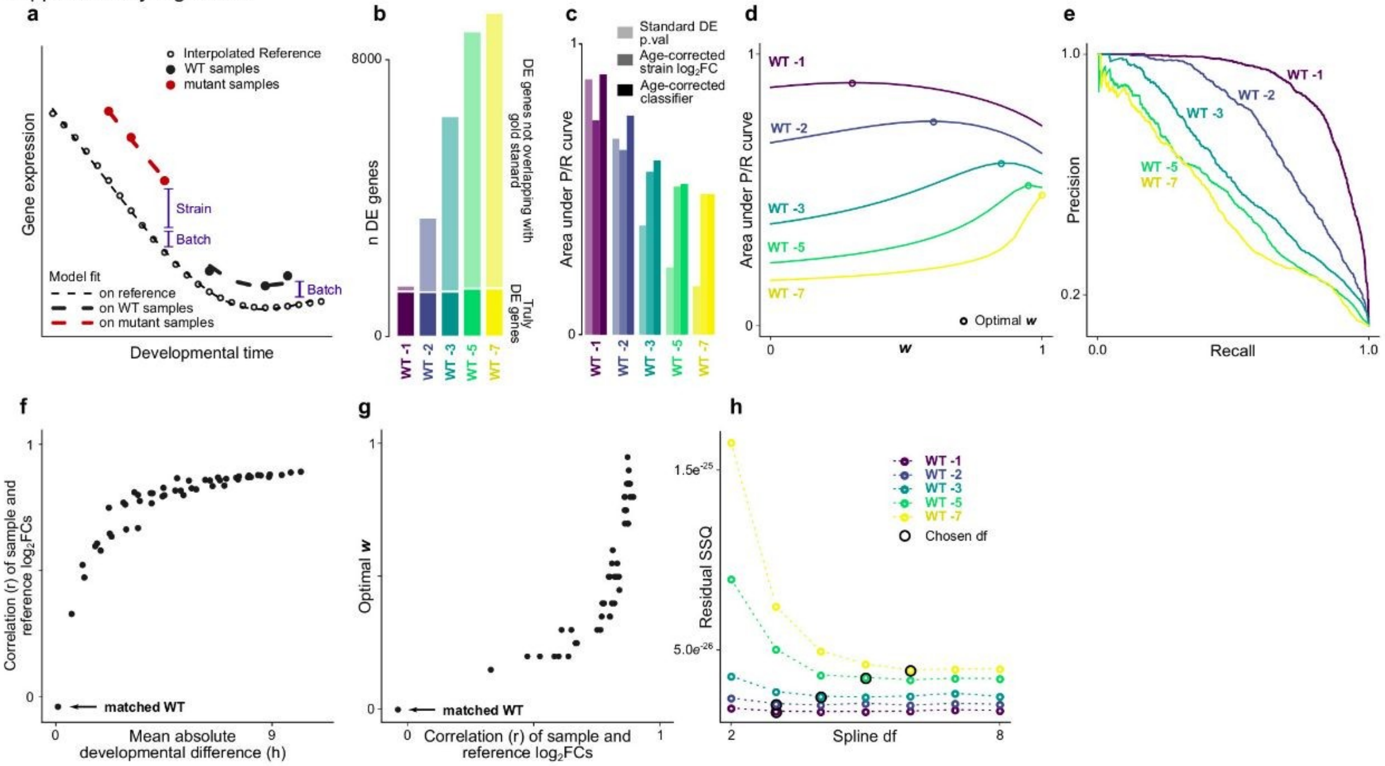


**Supplementary Figure 24 – Correcting the effect of development by integrating reference data**

Samples from *C. elegans* time-course experiments of WT and *xrn-2* mutants, profiled by Miki et al.[20] , and staged on the larval to young-adult reference built from Meeuse et al. samples[15] , are used to validate developmental correction approach. See also Fig 5e-h.

**a**, Cartoon of a model integrating a window of reference data, with Strain and Batch coefficients shown in blue.
**b**, Number of DE genes found by a standard DE model (FDR < 0.05) increases with the age gaps between compared groups, with a quasi-constant fraction of truly DE genes.
**c**, Area under precision-recall curves (AUPRC) in detecting gold-standard DE genes for standard DE model p-value, age-corrected logFCs, or the age-corrected classifier for each shifted WT subset.
**d**, w parameter optimization for shifted WT sets, by maximizing area under the Precision-Recall curves.
**e**, Precision-Recall curves of gold-standard gene detection by the age-corrected classifier for each shifted WT subset.
**f**, Correlation of expected development logFCs and observed logFCs between the *xrn-2* subset and combinations of 3-sample WT sets (note these are not the "WT -n" subsets, see Sup. Table 8).
**g**, Relationship between optimal w and sample-reference logFC correlation, as in (**f**).
**h**, Optimal spline degree-of-freedom (df) selection for the different WT shifted sets by reaching a residual Sum of Square (SSQ) plateau. The selected df increases with the shift, which is expected since the reference window to include gets larger and may thus contain more complex dynamics.

WT, wild-type. DE, Differential Expression or Differentially Expressed. logFC, log2 fold-change. FDR, false discovery rate.

#### References (Supplementary)


1. Aeschimann, F. *et al.* LIN41 post-transcriptionally silences mRNAs by two distinct and position-dependent mechanisms. *Mol. Cell* **65**, 476–489 (2017).
2. Alter, O., Brown, P. O. & Botstein, D. Singular value decomposition for genome-wide expression data processing and modeling. *Proc. Natl. Acad. Sci.* **97**, 10101–10106 (2000).
3. Storey, J. D., Xiao, W., Leek, J. T., Tompkins, R. G. & Davis, R. W. Significance analysis of time course microarray experiments. *Proc. Natl. Acad. Sci.* **102**, 12837–12842 (2005).
4. Reinke, V., San Gil, I., Ward, S. & Kazmer, K. Genome-wide germline-enriched and sex-biased expression profiles in Caenorhabditis elegans. *Development* **131**, 311–323 (2004).
5. Hendriks, G.-J., Gaidatzis, D., Aeschimann, F. & Großhans, H. Extensive oscillatory gene expression during C. elegans larval development. *Mol. Cell* **53**, 380–392 (2014).
6. Levin, M. *et al.* The mid-developmental transition and the evolution of animal body plans. *Nature* **531**, 637 (2016).
7. Collins, J. E. *et al.* Common and distinct transcriptional signatures of mammalian embryonic lethality. *Nat. Commun.* **10**, 1–16 (2019).
8. Rauwerda, H. *et al.* Transcriptome dynamics in early zebrafish embryogenesis determined by high-resolution time course analysis of 180 successive, individual zebrafish embryos. *BMC Genomics* **18**, 1–15 (2017).
9. Kim, D. hyun, Grün, D. & van Oudenaarden, A. Dampening of expression oscillations by synchronous regulation of a microRNA and its target. *Nat. Genet.* **45**, 1337–1344 (2013).
10. Domazet-Lošo, T. & Tautz, D. A phylogenetically based transcriptome age index mirrors ontogenetic divergence patterns. *Nature* **468**, 815–818 (2010).
11. Bahar, R. *et al.* Increased cell-to-cell variation in gene expression in ageing mouse heart. *Nature* **441**, 1011–1014 (2006).
12. Wang, X. *et al.* Gene expression patterns in the hippocampus during the development and aging of Glud1 (Glutamate Dehydrogenase 1) transgenic and wild type mice. *BMC Neurosci.* **15**, 1–17 (2014).
13. Chen, C.-Y. *et al.* Effects of aging on circadian patterns of gene expression in the human prefrontal cortex. *Proc. Natl. Acad. Sci.* **113**, 206–211 (2016).
14. Rockman, M. V., Skrovanek, S. S. & Kruglyak, L. Selection at linked sites shapes heritable phenotypic

variation in C. elegans. *Science* **330**, 372–376 (2010).
